# Entry and exit of *Fragilariopsis cylindrus* diatoms from the polar night: evidence for chromatin-mediated genome control

**DOI:** 10.1101/2025.04.02.646770

**Authors:** Nathalie Joli, Clara Bourbousse, Basile Leduque, Leïla Tirichine, Juliette Laude, Florian Maumus, Ouardia Ait-Mohamed, Louison Klein, Bran Kusendo, Lorenzo Concia, Leandro Quadrana, Fredy Barneche, Marcel Babin, Chris Bowler

## Abstract

Regulation of the epigenome landscape plays a crucial role in adaptation to environmental changes. However, the molecular mechanisms by which epigenome landscapes are established and tuned to adjust the cellular status to environmental cues remain poorly understood, especially in unicellular photosynthetic eukaryotes. Polar microalgae must adapt to extreme variations in light and temperature for survival and therefore represent excellent models to understand the diversity of gene regulatory mechanisms, yet chromatin-based mechanisms have never been explored in these organisms. In this study, we combined genome-wide profiling of histone and DNA modifications with transcriptomic analyses of the marine polar diatom *Fragilariopsis cylindrus* upon entry and exit of a 3-month-long polar night. We show that, in agreement with its established role in transcription facilitation, histone H2B monobiquitination (H2Bub) is modulated at active genes upon prolonged darkness to light re-exposure. In contrast, histone H3 lysine 27 trimethylation (H3K27me3) and DNA cytosine methylation (CG methylation), two modifications typically associated with the silencing of genes or transposable elements (TEs), remained stable throughout the transitions, reinforcing their role in transcriptional repression, particularly at TEs but also at a few genes. We further demonstrate that H3K27me3 and DNA methylation are physically associated at TEs, probably reflecting a dual locking system enabling *F.cylindrus* cells to cope with the invasive nature of these genetic elements under extreme environmental conditions. These findings enhance our understanding of genome and epigenome regulation in diatoms in response to environmental variability and open new avenues for exploring the role played by chromatin-level regulation in the unique evolutionary history of polar unicellular eukaryotes.

## Introduction

Diatoms, a diverse group of stramenopile microalgae, thrive in marine and freshwater ecosystems worldwide (Pierrella Karlusich et al., 2020). Although they make up less than 1% of the Earth’s biomass, diatoms contribute to ∼20% of global carbon fixation (Field et al., 1998). As shown with the *Phaeodactylum tricornutum* species, the exploration of genomic, metabolic and cellular features of diatoms provide insights into the ecological success of photosynthetic microorganisms in marine ecosystems (Falciatore et al., 2020) in which they form the basis of food webs, particularly in polar regions (Ibarbalz et al., 2019). Recent genome sequencing projects, including that of the polar diatom *Fragilariopsis cylindrus*, have started to shed light on the molecular underpinnings of diatoms’ extraordinary resilience to harsh conditions (Mock et al., 2017). Our previous study (Joli et al. 2024) identified that *F. cylindrus* survives the polar night by entering a quiescent state, where metabolic activity is drastically reduced, and cell division is arrested. This survival strategy is accompanied by internal structural changes that reflect the diatom’s morphological acclimation to darkness. Those observations underlined the remarkable robustness of polar diatoms, highlighting their ability to reprogram transcriptomic patterns to survive prolonged periods of darkness at low temperatures. Yet, mechanisms driving diatoms’ long-term transcriptional reprogramming remain poorly understood. In particular, the modulation of chromatin status may be at the nexus of signal integration and genome function during diatom quiescence and reactivation in response to environmental cues.

In eukaryotes, the modulation of genome organization into chromatin influences key cellular processes, including gene expression. Chromatin regulation is notably mediated by post-translational histone modifications and DNA methylation, which modulate chromatin structure, DNA accessibility, and transcriptional activity (Zentner et al., 2013; Law & Jacobsen, 2010). Chromatin mechanisms may thus play a pivotal role in the entry into and exit from quiescence in polar diatoms. Understanding how chromatin states mediate these adaptive responses is therefore crucial for unravelling the long-term survival strategies of polar diatoms.

While the definition of typical chromatin states associated to gene activation or silencing are largely concordant between plants and animals (Roudier et al., 2011; Sequeira-Mendes et al., 2014, Filion et al. 2010, Kharchenko et al., 2011), little is known in microalgae (Veluchamy et al. 2015). In this study we explore the epigenomic patterns of histone H2B mono-ubiquitination (H2Bub), a co-transcriptional modification conserved across yeast, plants, and mammals, which facilitates RNA polymerase II elongation and plays key roles in gene activation, tissue development, and disease (Tanny et al. 2007, Fuchs et al. 2014, Oss-Ronen et al. 2022). In the plant species *Arabidopsis thaliana*, H2Bub marks light-responsive genes within the first hour of illumination post-germination, enabling their efficient induction (Bourbousse et al. 2012), while its genome-wide levels and RNA Pol II transcriptional activity are globally reduced in darkness (Bourbousse et al. 2015, Nassrallah et al. 2018). The function of H2Bub in diatoms has never been assessed, yet, as in other organisms, it may be tightly linked to the transcriptional regime. We also explore potential variations in histone H3 trimethylation at lysine 27 (H3K27me3), a chromatin hallmark of gene repression during cell differentiation and development in animals and plants (Boyer et al. 2006, Schubert et al. 2005). In some, but not all eukaryotic species, H3K27me3 is deposited by Polycomb Repressive Complex 2 (PRC2), a chromatin modifier crucial for establishing and maintaining facultative heterochromatin. In contrast, constitutive heterochromatin is poorly accessible and typically marked by densely positioned DNA methylation and histone H3 lysine 9 methylation, altogether primarily silencing transposable elements (TEs). Interestingly, heterochromatinization can also influence nearby gene expression by creating a local environment refractory to transcription (Marsano et al. 2019). H3K27me3 and DNA/H3K9 methylation are therefore typically viewed as distinct silencing systems, yet this distinction has recently been questioned by evidence for a strong co-occurence of H3K27me3 and DNA methylation at TEs in certain eukaryotes (Hisanaga et al. 2023). Functional studies demonstrate that PRC2 activity plays a critical role in TEs silencing in unicellular microalgae and ciliates (Shaver et al. 2010, Frapporti et al., 2019, Zhao et al. 2019). In these organisms, H3K27me3 and in some cases, dual methylation of H3K9 and H3K27, is essential for repressing TEs. It has therefore been proposed that during the evolution of multicellular eukaryotes, PRC2 function was restricted to gene silencing while H3K9 methylation and DNA methylation specialized into silencing TEs and repeats (Deleris et al. 2021, Sharaf 2022).

Epigenomic studies in *P. tricornutum* have revealed both unique and conserved chromatin features in diatoms. Notably, chromatin marks as H3K4 dimethylation (H3K4me2), acetylation of H3K9 and H3K14 (H3K9/14Ac) and H3K9/DNA methylation were shown to vary at particular gene sets under upon exposure to nutrient-limiting conditions, demonstrating the dynamic nature of the diatom epigenome landscape in face of environmental variation (Veluchamy et al., 2015). DNA methylation and H3K27me3 marking have both been observed at TEs as well as a few genes, correlating with a low transcript readout (Veluchamy et al., 2013, 2015). Still in the in *P. tricornutum* model species, knocking out PRC2 activity disrupts cell morphology and thermomorphogenesis underscoring the role of this activity in diatom cell differentiation (Zhao et al., 2021, Zarif et al., 2024).

By exposing *F. cylindrus* to transitions between light and prolonged darkness mimicking the polar night, we explored the links between epigenome adaptations and the regulation of genes and TEs. Following a complete reannotation of *F. cylindrus* TEs, we observed that these genetic elements are dually marked by H3K27me3 and DNA methylation, a phenomenon previously inferred in other diatom species only through correlations. We also unravel the H2Bub genome-wide landscape in diatoms, which, as in multicellular eukaryotes, is associated with active transcription. Under simulated polar night followed by light re-exposure, H2Bub levels dynamically adjust, targeting distinct gene sets in response to light availability, thereby probably contributing to fine-tune gene expression involved in essential functions for survival. In contrast, the repression of genes and TEs marked by H3K27me3 and DNA methylation remains largely unaffected by light shifts, highlighting the stability of these epigenetic features upon extreme environmental changes. We propose that a dual-locking mechanism, combining PRC2-mediated repression and DNA methylation, ensures the repression of invasive genetic elements to safeguard genome integrity against profound environmental perturbations.

## Materials and methods

### Experimental set-up

*F. cylindrus* CCMP 3323 cells were cultured as described in Joli et al. (2024), with sampling time points detailed in **Figure 1**. Briefly, triplicate cultures were acclimated to full light conditions (30 μmol photons m² s⁻¹) in a cold room at 0 ± 1°C in 80 L glass cylinders. Following acclimation, cultures were transferred to total darkness in a closed system for three months. Subsequently, the cultures were returned to full-light conditions for seven days. Importantly, the samples subjected to three months of darkness were unintentionally exposed to light during transfer from the cylinders to the centrifuge during sampling. Consequently, this sample cannot be considered a true darkness sample but rather one briefly exposed to light (labelled RL5sec).

**Figure 1:**
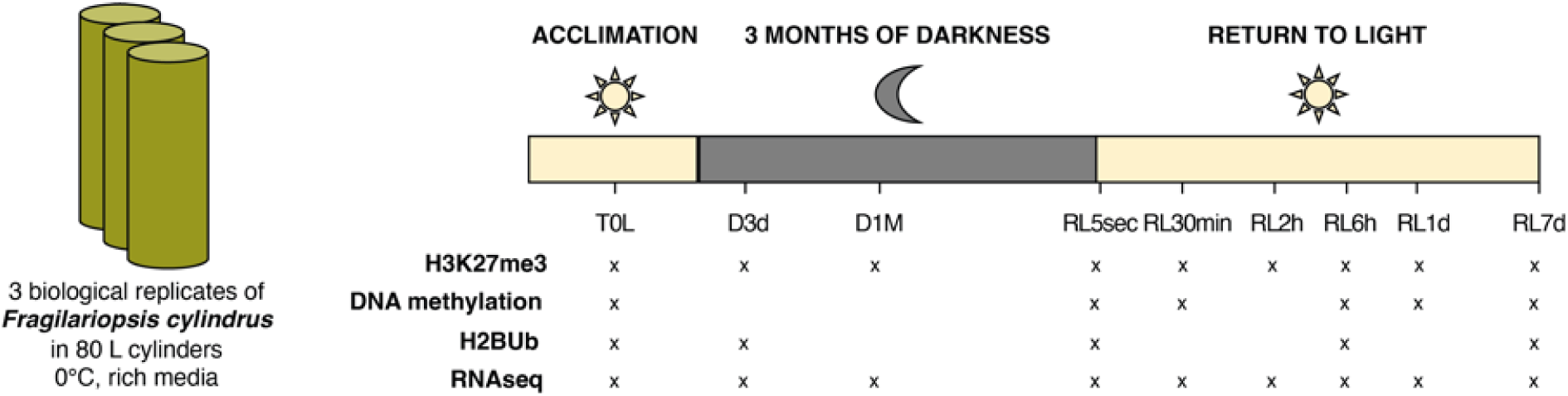
Experimental design and timing of transcriptomic and epigenomic measurements. The cylinders depicted here are the same as those described in Joli et al. (2023). T0L (for T0 light) represents the acclimation phase to full light, followed by 3 months of darkness (D for Dark) and the subsequent return to light period (RL for light return). The time series differ between transcriptomic and epigenomic measurements, prioritizing a comprehensive yet feasible experimental approach.

### Sample Collection

Samples for RNA and DNA extractions were collected and preserved following the protocol outlined in Joli et al. (2024). Cells were harvested by centrifugation, rinsed in marine PBS (mPBS) solution (490 mM NaCl, 15 mM Na2HPO4, 4.5 mM NaH2PO4, 0.1 mM EGTA), and the resulting pellets were flash-frozen and stored at -80°C until extraction.

For histone modification profiling (H2Bub and H3K27me3), cells were crosslinked immediately after harvesting. Crosslinking was performed by incubating cells in a 1% formaldehyde solution prepared in mPBS at 0 ± 1°C for 30 minutes. The reaction was quenched by adding glycine to a final concentration of 0.12 M and incubating for 15 minutes. Crosslinked cells were washed twice with cold mPBS and pelleted by centrifugation. The resulting dry pellets were flash-frozen in liquid nitrogen and stored at -80°C for downstream analyses.

### Chromatin extraction

Chromatin was extracted using a modified version of the method developed by Lin et al. (2012). Fixed frozen cells were resuspended in Extraction Buffer 1 (0.4 M sucrose, 10 mM Tris-HCl, pH 8.0, 10 mM MgCl₂, 5 mM 2-mercaptoethanol, 1% Triton X-100, supplemented with a Roche protease inhibitor cocktail, cat. COEDTAF-RO). To facilitate cell disruption and break the silica frustules, a bead-beating step was performed at 30 Hz for 5 minutes using a high-speed disruptor and a mixture of glass beads (equal parts of 212–300 µm and 425–600 µm beads; Sigma, cat. G1277 and G8772). The lysate was diluted with additional Extraction Buffer 1 and subjected to 20 minutes of chemical lysis, with brief vortexing (10 seconds every 5 minutes). To remove beads and cellular debris, the lysate was filtered through a double layer of Miracloth. Intact organelles from the flow-through were pelleted by centrifugation at 3,300 × g for 20 minutes. The resulting pellet underwent two washing steps, each consisting of resuspension in Extraction Buffer 2 (0.25 M sucrose, 10 mM Tris-HCl, pH 8.0, 10 mM MgCl₂, 5 mM 2-mercaptoethanol, 1% Triton X-100, protease inhibitors) and centrifugation at 9,000 × g for 10 minutes. The washed pellet, containing nuclei, was further processed by resuspension in Nuclei Lysis Buffer (50 mM Tris-HCl, pH 8.0, 10 mM EDTA, 1% SDS, protease inhibitors). Insoluble debris were removed by centrifugation at 16,000 × g for 5 minutes, and the chromatin-rich supernatant was flash-frozen in liquid nitrogen and stored at -80°C for further use. Throughout the procedure, all solutions and equipment were pre-chilled, and all steps were conducted on ice. Centrifugations were carried out at 4°C. Chromatin protein concentrations were measured using the Pierce™ BCA Protein Assay Kit.

### Chromatin immunoprecipitation (ChIP) analyses

H3K27me3 and H2Bub ChIPs were performed using 400 µg of chromatin following the protocol by Lin et al. (2012) with minor modifications. For each histone mark, two biological samples per time point were considered. Chromatin samples were sonicated using an ultrasound sonicator (Bioruptor®, Diagenode; 1x 15 minutes, High, 30 seconds ON – 30 seconds OFF), generating 200 to 700 bp DNA fragments. An aliquot was reserved as the total DNA sample or “INPUT”. Chromatin samples were diluted with ChIP Dilution Buffer (1% Triton X-100, 1.2 mM EDTA, 16.7 mM Tris-HCl, pH 8.0, 167 mM NaCl, 0.1 mM PMSF, MiniTAB EDTA-free) and divided into three tubes: two for immunoprecipitation (“IP”) and one for the no antibody control (“Mock”). For each IP, equal amounts of A beads and G beads were mixed (Invitrogen cat. No. 100.04D, (Invitrogen cat. No. 100.02D) and washed with ChIP Dilution Buffer. This suspension was then split into two aliquots. 5 µL of antibody (H3K27me3: C36B11, Cell Signaling; H2Bub: MM-029-P, Medimabs Millipore) was added to one of the aliquots of washed bead suspension for antigen capture. The remaining chromatin solution was added to the second aliquots of washed bead suspension for preclearing, and the tubes were rotated gently at 4°C for 3 hours. Following preclearing, the chromatin was transferred to the antibody-bound beads and incubated at 4°C overnight with gentle rotation. Following immunoprecipitation, bead-antibody-antigen complexes were washed four times using Low Salt Wash Buffer, High Salt Wash Buffer, LiCl Wash Buffer, and TE Buffer. After washing, the immune complexes were eluted twice from A/G beads by adding 250 µL of Elution Buffer and incubating at 65°C for 15 minutes with vortexing for mixing.

From here, INPUT samples were treated in parallel of IP and Mock eluates. Crosslinking was reversed by the addition of NaCl (final concentration 200 mM) and incubation at 65°C overnight. After crosslink reversal, the samples were incubated at 45°C for 1 hour with Proteinase K to degrade proteins. DNA was then extracted using phenol-chloroform-isoamyl alcohol, followed by precipitation with ethanol, 1/10 volume of 3 M sodium acetate (pH 5.3), and GlycoBlue™ (Invitrogen, cat. AM9515). Samples were incubated at -80°C overnight to ease DNA precipitation. Following centrifugation at 16,000 x g for 1 hour at 4°C, the DNA pellet was washed with 70% ethanol. The residual liquid was removed, and the pellet was air-dried under a fume hood using a 55°C dry bath. Finally, DNA pellets were resuspended in sterile molecular-grade ddH2O. ChIP sequencing libraries were constructed and sequenced at the Fasteris facility using the NextSeq platform (single-end, 1 x 50 bp). The sequencing output averaged 15 million clean reads per sample for INPUT controls and 30 million reads per sample for IPs.

ChIP bioinformatic analysis followed the publicly available pipeline (https://github.com/vidal-adrien/ChIP-Rx-Pipeline-Pub). Raw sequencing data quality was assessed with FastQC v0.11.9. The trimmed reads (Trimmomatic v0.38) were aligned to the *F. cylindrus* CCMP1102 reference genome (Mock et al., 2017) using STAR v2.7.9a. Peak calling was performed with MACS2 v2.2.6 callpeak, applying custom parameters for H3K27me3 ChIP, with the mfold parameter set to a range of 3 to 30, to optimize peak detection while minimizing false positives. Peak annotation over genes and TEs was performed using bedTools v2.27.1. Heatmaps and metaplots were generated with deepTools (computeMatrix, plotProfile, plotHeatmap). Differentially marked genes were identified using the DESeq2 R package v1.44.0.

For H2Bub ChIP, because distinct peaks were not discernible in one of the biological replicates for two time points (due to elevated background noise in certain tracks), we decided to focus on analyzing H2Bub enrichment over genes using the immunoprecipitation (IP) signal. This approach was justified by our observation that, when examining the global gene dataset, the IP signal over genes consistently surpassed the input signal.

### Transcriptomic analyses

RNA was isolated as described in Joli et al. (2024), using the RNeasy Mini Kit (Qiagen) following the protocol of the manufacturer. A second DNase step was added after elution (DNase Invitrogen™ Kit TURBO DNAfree). RNA libraries were prepared using the Illumina TruSeq Stranded mRNA Library Prep Kit, which includes poly-A selection. Sequencing was performed by Fasteris using the Illumina HiSeq 4000 platform, generating >30 million clean reads per sample (paired-end, 2 × 150 bp). RNA deep sequencing raw data were previously deposited at https://www.ncbi.nlm.nih.gov/geo/ under the GEO accession no. GSE218215 (Joli et al., 2024). Read quality was assessed using FASTQC (v0.11.9). The trimmed reads (Trimmomatic v0.38) were aligned to the reference genome using STAR v2.7.9a. Raw read counts using TEtranscripts (Jin et al., 2015) and used for differential expression analysis with DESeq2 v1.26.0 which accounts for ambiguously mapped reads between genes and transposable elements using the acclimation phase to full light before darkness (T0L) as the reference time point.

### Methylome analyses

Genomic DNA was extracted following the protocol #3 of the Easy-DNA Kit (Invitrogen Life Technologies, Carlsbad, CA, USA). Whole-genome bisulfite sequencing (BS-seq) libraries were constructed and sequenced at the BGI Genomics Hong Kong facility using the BGI Genomics pipeline and DNBSEQ™ platform (MGISEQ-2000), yielding >20 million clean reads per sample (paired-end, 2 × 100 bp). Paired-end WGBS reads were trimmed using Trimmomatic program (version 0.33) with parameters “ILLUMINACLIP:TruSeq3-PE.fa:2:30:10 LEADING:3 TRAILING:3 SLIDINGWINDOW:4:15 MINLEN:36” (Bolger, Lohse, and Usadel 2014). Mapping of trimmed sequences to *F. cylindrus* reference genome with option “-n 1 -l 20”, removal of identical reads, and counting of methylated and unmethylated cytosines were performed by Bismark ver. 0.15.0 (Krueger and Andrews 2011). MethylKit package v0.9.4 (Akalin et al. 2012) was used to calculate differential CG methylation in 100 bp non-overlapping windows (DMRs) between the different time points and T0L. Significance of calculated differences was determined using Fisher’s exact test and Benjamin-Hochberg (BH) adjustment of p-values (FDR<0.05) and methylation difference cutoffs of 40%.

To assess potential co-marking of H3K27me3 and DNA methylation in the same nucleosomes, H3K27me3 ChIP DNA from all light and dark samples were merged to reach 144ng total DNA. INPUT DNA was also merged to obtain over 1ug DNA. ONT libraries (SQK-*LSK110*) were prepared using the ChIP and INPUT DNA and loaded into R9.4.1 Minion flow-cell. Sequencing was performed in a Mk1C device during 72hs. Basecalling was performed using Guppy with the HAC model, yielding 220,000 and 516,695 reads with an average size of 471nt and 298nt for IP and INPUT DNA, respectively. ONT reads were mapped to the reference *F. cylindrus* genome using minimap2 with default parameters. mCG methylation calling was performed using DeepSignal-plant (Ni et al. 2021).

### Annotation of the transposable elements

A de novo repeat detection analysis was performed with the TEdenovo pipeline from the REPET package (v2.4; Flutre et al., 2011) to generate a library of consensus sequences. Whole-genome annotation was then conducted using TEannot (Quesneville et al., 2005; Dataset S2), utilizing this repeat library as a digital probe.

Repeats were found to represent approximately 18% of the *F. cylindrus* 60 Mb genome, encompassing various categories of TEs, simple sequence repeats (SSRs), and satellite sequences (SAT). For our analysis, we focused exclusively on well-defined TE categories of minimal size of 20 bp, including Gypsy, Copia, DIRS, LINE, PiggyBac, Tase, TIR MITE, and Crypton. This selection resulted in a final dataset of 8,096 TEs for further exploration. The resulting repeat annotations were combined with the *F. cylindrus* reference genome (Mock et al., 2017) creating a comprehensive dataset of gene and TEs features.

### Functional enrichment

Gene Ontology (GO) term enrichment analysis was performed using the topGO R package to identify over-represented GO categories in the set of differentially expressed genes. The Over-Representation Analysis (ORA) method was applied to test whether specific GO terms were significantly enriched among the genes of interest compared to a background set. Fisher’s exact test was used to determine statistical significance, with a threshold of P < 0.05. The results were summarized to a shorter list using Revigo (Supek et al. 2011).

### Identification of putative homologs

To identify putative orthologs involved in the deposition of the three chromatin marks of interest (H2Bub, H3K27me3, and CG DNA methylation), we first searched for homologs using Kegg Orthology (KO) annotation, which assigns genes to conserved functions based on standardized identifiers. For candidates lacking a KO number, such as the E(Z) subunit of the PRC2 complex, we performed BLASTp searches against multiple species, including *P. tricornutum*, followed by a BLASTp search against the non-redundant (nr) database to validate the identified candidates. Additionally, we refined our dataset using a phylogeny-based classification of DNMTs (Hoguin et al., 2023), allowing us to include the different DNMT families previously described. Finally, for DNMT candidates with a KO number but absent from this classification, multiple sequence alignments were performed using MAFFT in Geneious. The neighbor-joining tree, built in Geneious, includes all DNMTs from *P. tricornutum* and *F. cylindrus* to identify their closest homologs.

### Transcription factor binding site identification

Transcription factor (TF) binding sites within the TEs sequences of interest were identified using tfscan, a tool from the EMBOSS suite (Rice et al., 2000). This tool scans DNA sequences for potential TFBS based on the TRANSFAC database. Binding site matches were detected using a fast sequence word-matching algorithm, with no mismatches allowed.

## Results

### H2Bub levels at genes correlate with their expression levels over time in darkness and upon re-exposure to light

*F. cylindrus* undergoes significant gene expression reprogramming when subjected to prolonged darkness followed by a return to light (Joli et al., 2024). To explore the role of chromatin regulation in this process, we first focused on the co-transcriptional H2Bub mark. We identified two putative homologs of *Bre1*, the E3 ubiquitin ligase catalyzing H2B monoubiquitination in yeast. Notably, both homologs exhibited higher expression levels under light conditions compared to prolonged darkness (**Table S1**). Although previously validated in *A. thaliana* (Roudier et al., 2011), the H2Bub antibody used in chromatin immunoprecipitation assays here yielded suboptimal results, yet it enabled to define global trends at gene sets. Indeed, ChIP-seq profiling of H2Bub in *F. cylindrus* revealed a distribution pattern consistent with other species, with an enrichment primarily over gene bodies. Analysis of H2Bub levels at marked genes for each condition further indicate a reduction in H2Bub levels after three months of complete darkness minimally exposed to 5 seconds of light (RL5sec) compared to all the other time points (**Figure 2A**).

**Figure 2:**
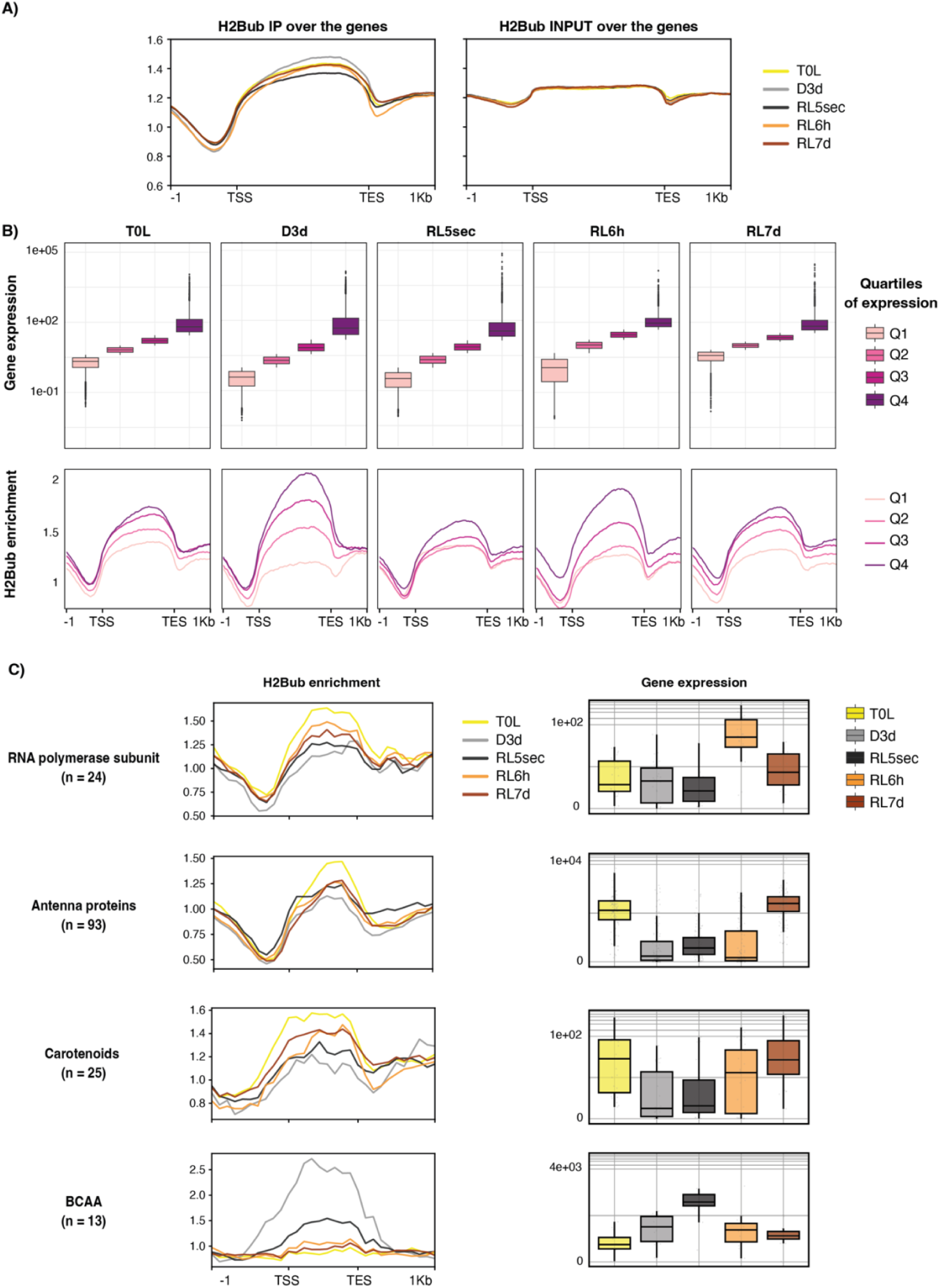
H2Bub abundance over gene bodies correlates with expression levels and targets distinct gene sets associated with different functions in response to light availability. **(A)** Average H2Bub occupancy across all genes in light-to-dark-to-light conditions in the IP and INPUT samples. **(B)** Boxplot of genes with different expression quartiles based on normalized read counts at each time point (Q1: top 25% least expressed genes, Q2: between the 25th and 50th percentile, Q3: between the 50^th^ and 75th percentile, Q4: top 25% most highly expressed genes) and corresponding metaplots of H2BUb enrichment (IP) across the same genes. **(C)** H2Bub enrichment at genes associated with RNA polymerase subunits, light-harvesting antenna proteins, carotenoid biosynthesis, and the branched-chain amino acid pathway, compared to gene expression over time (number of genes in parentheses). Gene lists were extracted from Joli et al. (2024). The RL5sec sample corresponds to cells exposed to three months of darkness followed by a 5-second flash of light.

Given the well-established link between H2Bub and transcriptional activation in yeast, animal, and plants (Henry et al. 2003, Minsky et al. 2008, Grasser et al. 2021), we examined its relationship with gene expression. Genes were grouped into quartiles based on expression levels, revealing a clear positive association: genes with the highest expression at each time point showed the strongest H2Bub enrichment, while those with lower expression exhibited reduced H2Bub levels (**Figure 2B**). We also observe that while H2Bub mean enrichment was highly different between gene sets with distinct expression levels during transitions from continuous light to darkness (D3d) and from darkness to light (RL6h), this dynamic is much weaker at the RL5sec time point. Considering this time point as a dark-adapted state, this observation aligns with the hypometabolic state of *F. cylindrus* under prolonged darkness during which genes tend to be poorly expressed (**Figure 2B**). Notably, the composition of gene sets within the expression quartiles shifted over time (**Figure S2)**, reflecting the extensive transcriptional reprogramming observed during prolonged darkness and light recovery (Joli et al., 2024).

We further examined H2Bub enrichment over genes involved in functions essential for survival in darkness and transition to light (**Figure 2C**). Genes encoding RNA polymerase subunits, light harvesting antenna proteins, and carotenoid biosynthesis enzymes show higher H2Bub enrichment and increased expression in light conditions, consistent with the biogenesis of the photosynthetic apparatus involving transcription reactivation. In contrast, genes involved in the branched-chain amino acid (BCAA) pathway exhibit greater H2Bub enrichment and mRNA levels during darkness. Both the changes in amplitude of H2Bub enrichment at gene sets with distinct expression ranges, and the differential marking of light vs dark-expressed key genes, highlight the dynamic regulation of H2Bub. Altogether these observations highlight a profound adaptation of the epigenome landscape at regulated genes during *F. cylindrus* polar night entry and exit.

### H3K27me3 is predominantly located at TEs but also at a few genes

To assess H3K27me3 deposition and function in *F. cylindrus*, we first searched for putative homologs of the Polycomb Repressive Complex 2 (PRC2) complex subunits. PRC2 consists of four conserved subunits across eukaryotes: the histone methyltransferase E(Z), the WD40 domain protein ESC, the zinc finger protein SUZ12, and the nucleosome remodeling factor NURF55 (Chittock et al., 2017; Schuettengruber et al., 2017; Dumesic et al. 2015). We identified clear candidates for E(Z), ESC, and NURF55, but not for SUZ12 (**Table S1**). These genes are generally more expressed upon light re-exposure after darkness. We next examined the genome-wide distribution of the H3K27me3 mark by ChIP-seq. The distribution of H3K27me3 exhibits a broad pattern with large peaks (**Figure S3**), reminiscent of its distribution in animals (Margueron and Reinberg, 2011). H3K27me3 altogether covers approximately 9 Mb, i.e. 15% of the 60 Mb-long assembled *F. cylindrus* genome, a coverage comparable to the model diatom *P. tricornutum* (Veluchamy et al., 2015) and approximately twice more than in *A. thaliana* (Roudier et al., 2011). H3K27me3 peaks were detected in a small subset of gene bodies, representing 5% of the total annotated genes across all time points. In contrast, the mark was predominantly associated with TEs, with 36% of annotated TEs showing significant marking (**Table 1**), a pattern consistent with previous findings in *P. tricornutum* (Veluchamy et al., 2015).

**Table 1:** Predominant association of H3K27me3 marks with TEs over genes, covering at least 80% of the coding sequence at one or more time points.

|  | **GENES** | **TEs** |
| --- | --- | --- |
| Total number | 27140 | 8096 |
| Marked by H3K27me3 at least once | 1338 (5%) | 2890 (36%) |

To get a deeper understanding of TEs chromatin status, we undertook a complete reannotation of TEs in the *F. cylindrus* genome. The majority of TEs identified in *F. cylindrus* are Class 1 TEs that replicate through reverse transcription of an RNA intermediate, as opposed to Class 2 TEs that use a “cut and paste” mechanism, as observed in the *P*. *tricornutum* and *Thalassiosira pseudonana* genomes (Maumus et al. 2009). A similar distribution is observed among the H3K27me3-marked TEs, which are predominantly classified as Class 1, with a slight enrichment in *Copia* elements (**Figure S5**).

A large proportion of the H3K27me3-marked genes has unknown functions. Yet, for 314 of these 1338 genes (**Table 1**), we observed an enrichment in key biological processes such as chromatin remodeling, carbohydrate transport for energy metabolism, regulation of protein modifications essential for signaling, cellular stress responses (including DNA repair), and ubiquitin-protein transferase activity associated with protein degradation and turnover. Notably, this subset includes genes encoding heterochromatin-associated proteins, including Heterochromatin Protein-1 (HP1), a chromatin-associated protein that physically associate with H3K9me2/3 to promote heterochromatin formation, and CENP-B, a protein required for the incorporation of the centromere-specific CENP-A histone variant, as well as components of chromatin remodeling complexes and bZIP-type transcription factors (TF) (**Table S2**). Of note, genes enriched with the H2Bub mark and actively expressed under light or dark conditions tend to be depleted of H3K27me3 (**<u>Figure S4</u>**). This observation suggests that the two histone modifications occupy distinct chromatin compartments and may have an antagonistic relationship in *F. cylindrus*.

### Despite slight attenuations in darkness, H3K27me3 marking and repression of genes TEs are maintained during prolonged darkness

We investigated the temporal dynamics of H3K27me3 patterns at the T0L, D3d, D1M, RL5sec, RL30min, RL2h, RL6h, RL1d and RL7d time points. This showed a global decline in H3K27me3 enrichment in darkness, followed by a progressive restoration upon light exposure at both genes and TEs seven days after the return to light (**Figures S6A, S6B**). Despite this attenuation in darkness, genes and TEs marked by H3K27me3 maintain a high enrichment at all time points as compared to unmarked genes and TEs (**Figures S6C, S6D**). We further investigated changes in H3K27me3 enrichment over time by performing a differential analysis of H3K27me3 peaks. This analysis revealed negligible changes (|log2FC| < 0.5) between time points, indicating that H3K27me3 levels remain largely stable despite a slight overall decrease observed under prolonged darkness. Building on this observation, we next examined the expression dynamics of these H3K27me3-marked elements.

Genes marked by H3K27me3 across their entire bodies expectedly exhibit notably low transcript levels, with only 3% showing any detectable RNA-seq reads (normalized read counts > 10) (**Figure 3A, B**). In contrast, genes lacking the H3K27me3 mark display higher expression frequencies (76%) and significantly higher expression levels (**Figure 3A, B, C).** Interestingly, a subset of genes displays H3K27me3 marking localized exclusively to their 5’ or 3’ regions (**Figure 3**). These genes are more frequently expressed compared to those with the mark spread across the entire gene body, suggesting partial transcriptional repression when the mark is restricted to the gene’s borders. Notably, genes marked in 5’ regions, which plausibly correspond to promoter proximal domains, exhibit significantly lower expression levels than unmarked genes. After examining the genomic context of all genes marked exclusively in their 5’ proximal region, we found them to be enriched in TEs within the upstream 2 kb flanking region. Similarly, genes marked in the 3’ region are associated with a higher density of TEs in the downstream flanking regions (**Table S3**). This suggests a potential influence of heterochromatic TEs on some neighboring genes, H3K27me3 broad domains plausibly spreading from TEs and creating a chromatin state refractory to gene transcription.

**Figure 3:**
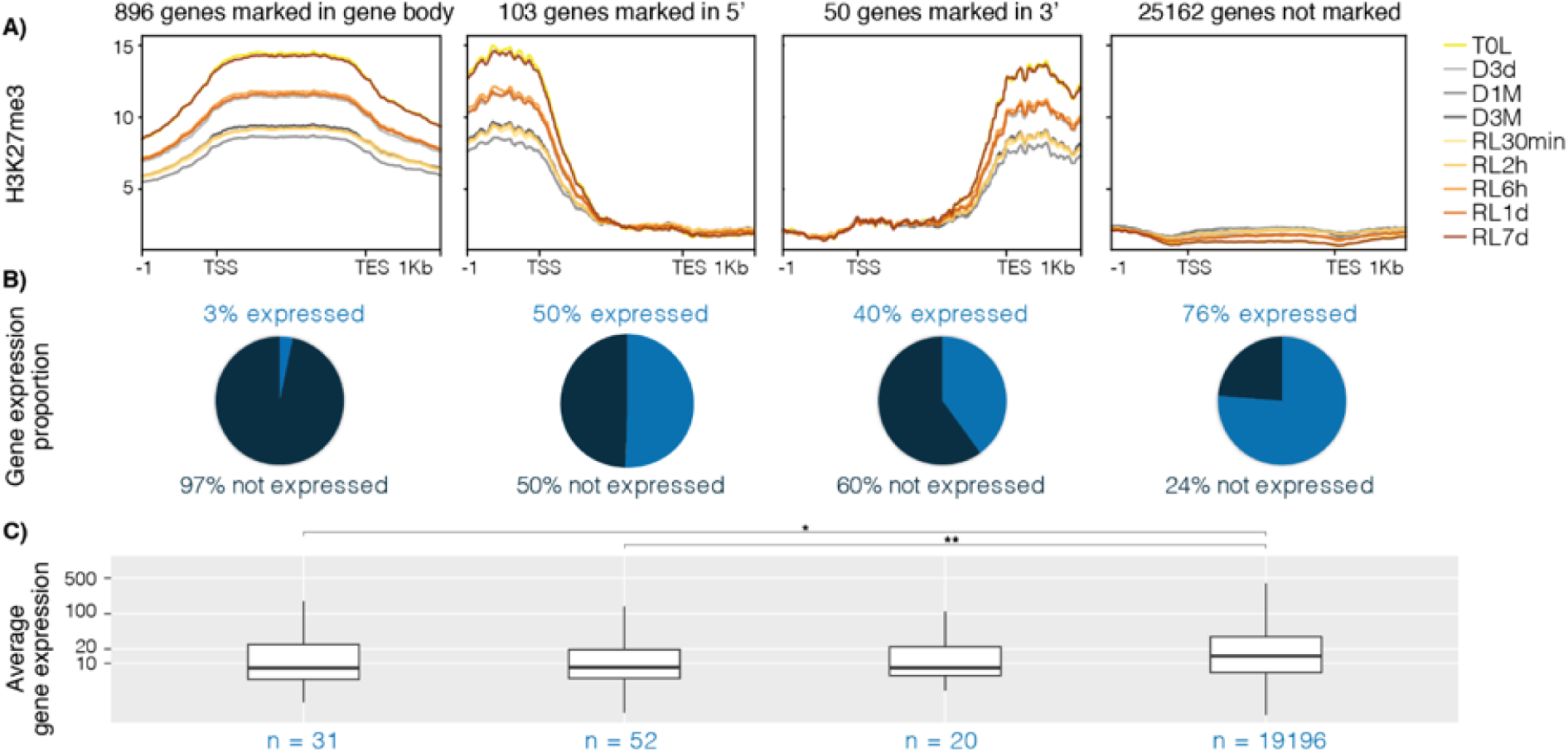
H3K27me3-marked genes are poorly expressed. (**A**) Metaplot of H3K27me3 enrichment across genes marked at the gene body, at the 5’ end, at the 3’ end, and unmarked genes. (**B**) The associated proportion of expressed genes. **(C)** Average read counts for the corresponding gene sets. The statistical analysis includes Mann-Whitney U tests and pairwise Wilcoxon tests with Holm correction, with significance levels indicated as follows: p < 0.05 (*), p < 0.01 (**).

Like genes, TEs marked by H3K27me3 tend to show lower expression levels than unmarked TEs, with less than 1% showing any detectable RNA-seq reads (normalized read counts > 10) (**Figure 4**). These findings align with the well-established role of H3K27me3 as a repressive mark not only when located at gene bodies as in fungi, plants, and metazoans (Jamieson et al., 2013; Zhang et al., 2007; Pauler et al., 2009), but also at TEs, potentially influencing genome stability.

**Figure 4:**
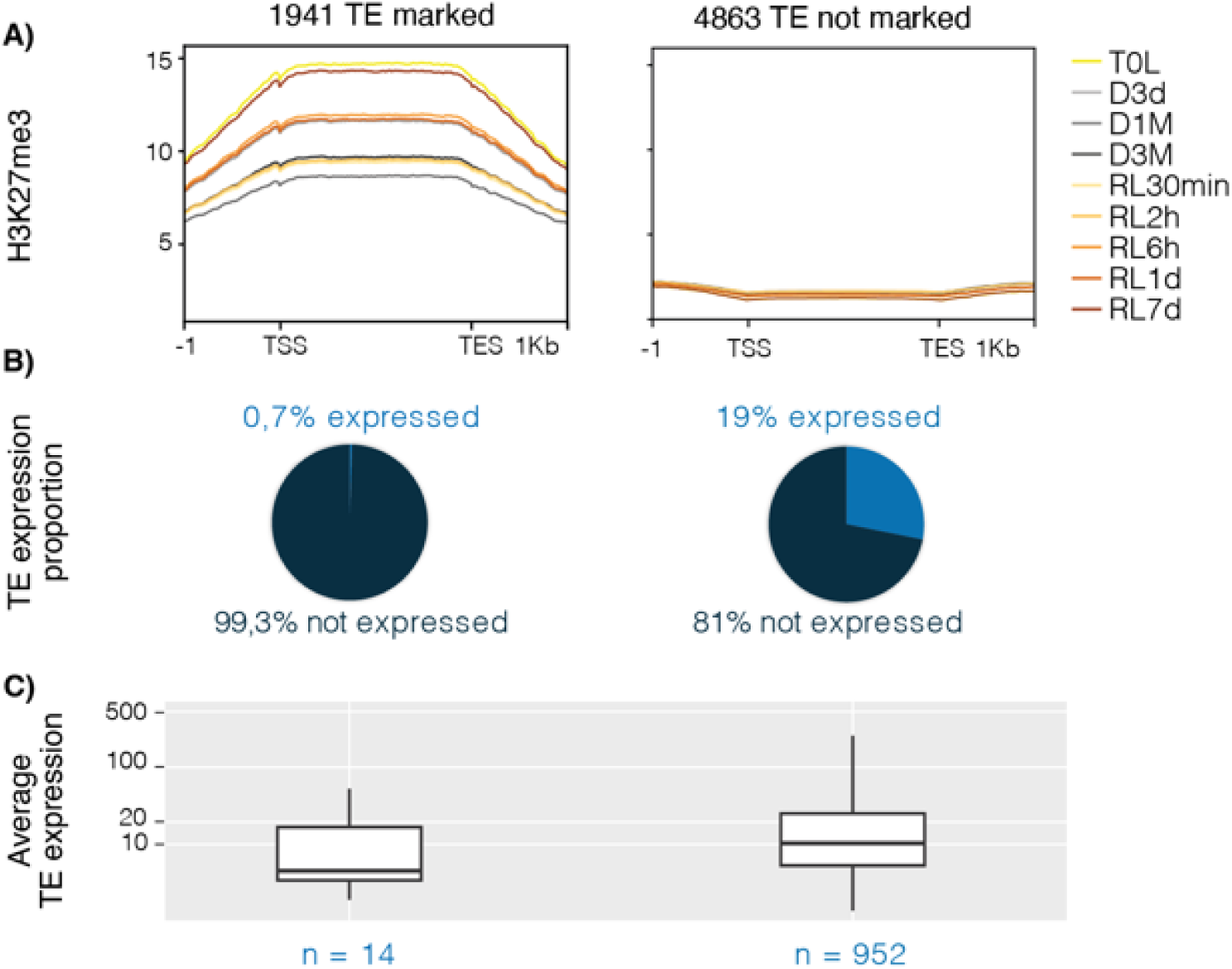
H3K27me3 associates with the extent ofTE silencing in *F. cylindrus*. (**A**) Metaplot of H3K27me3 enrichment across H3K27me3-marked and unmarked TEs. (**B**) Proportion of expressed TEs. **(C)** Average read counts at H3K27me3-marked and unmarked TEs.

All our attempts to identify potential correlations between H3K27me3 enrichment and the expression of the few H3K27me3-marked genes and TEs with detectable read counts failed to reveal any significant relationship.

We identified two particularly interesting TEs based on their genomic context. These elements are marked by H3K27me3, are differentially expressed in response to prolonged darkness and located within intronic regions overlapping gene annotations (**Figure S7**). Both belong to Class 1 long terminal repeats (LTRs) of the Copia and Gypsy families. Additionally, we identified TF binding sites within their sequences (**Table S4**). These observations suggest that they may play a role in modulating transcription, potentially through interactions with TFs.

### DNA methylation remains stable throughout light and dark conditions

DNA methylation plays a critical role in regulating various biological processes, including gene and TE silencing in plants (Kato et al., 2003). In the reference genome of *F. cylindrus*, we identified eight putative DNMT homologs, encoding DNMT4, DNMT5, and DNMT6 enzymes (**Table S1**). One of the DNMT4-like candidates is expressed after one month of darkness while the others show stronger expression when light returns. Using BS-seq, we generated whole-genome methylome data for *F. cylindrus*. This identified DNA methylation across 4.3 Mb, coresponding to 7.2% of the 60 Mb genome, a higher density than in the 5% estimated for *P. tricornutum* genome (Veluchamy et al., 2013).

Methylation was significantly detected in CG, CHG, and CHH contexts (where H represents any nucleotide except G), with over 80% occurring in the CG context (**Figure S8**).

To explore CG methylation dynamics, we conducted a differential analysis to identify genes and TEs with methylation changes in response to prolonged darkness and light return, T0L used as reference condition. Our results revealed a small number of differentially methylated regions (DMRs): 15 hypermethylated genes, 14 hypomethylated genes, 18 hypermethylated TEs, and 23 hypomethylated TEs, with nearly no overlap between time points (**Figure S9)**. Further analysis of the methylation profiles, compared with gene expression and H3K27me3 patterns, showed that the few genes and TEs identified as being differentially methylated are not expressed. Overall, these analyses showed that the *F. cylindrus* methylome is extremely stable under prolonged darkness and light return while genes and TEs are also stably repressed.

### DNA associated with nucleosomes marked by H3K27me3 is methylated

Building on previous observations that H3K27me3 and DNA methylation often correlate over TEs and genes in certain unicellular organisms, we examined the distribution of these marks across all genes and TEs (**Figure 5**). Our analysis reveals strikingly similar patterns, with both chromatin marks aligning in nearly identical genomic regions, whether on genes or TEs. Furthermore, this colocalization of H3K27me3 and DNA methylation is associated with a lack of expression of both genes and TEs. To further investigate this relationship, we mapped DNA methylation tracks to genes and TEs classified as either marked or unmarked by H3K27me3. The results revealed that DNA methylation largely coincides with regions marked by H3K27me3, and both remain stable and associated across all time points (**Figure S10A**).

**Figure 5:**
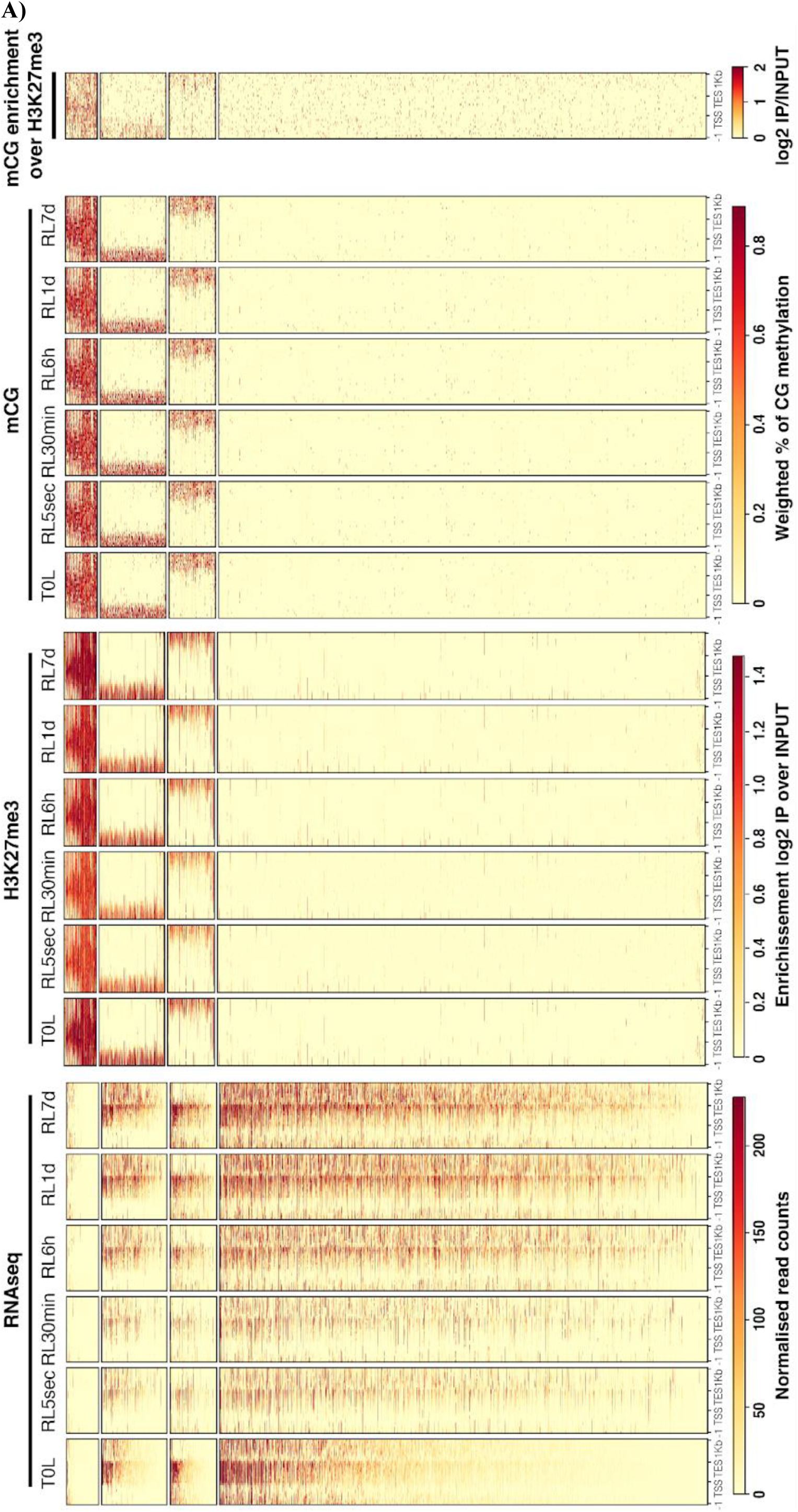
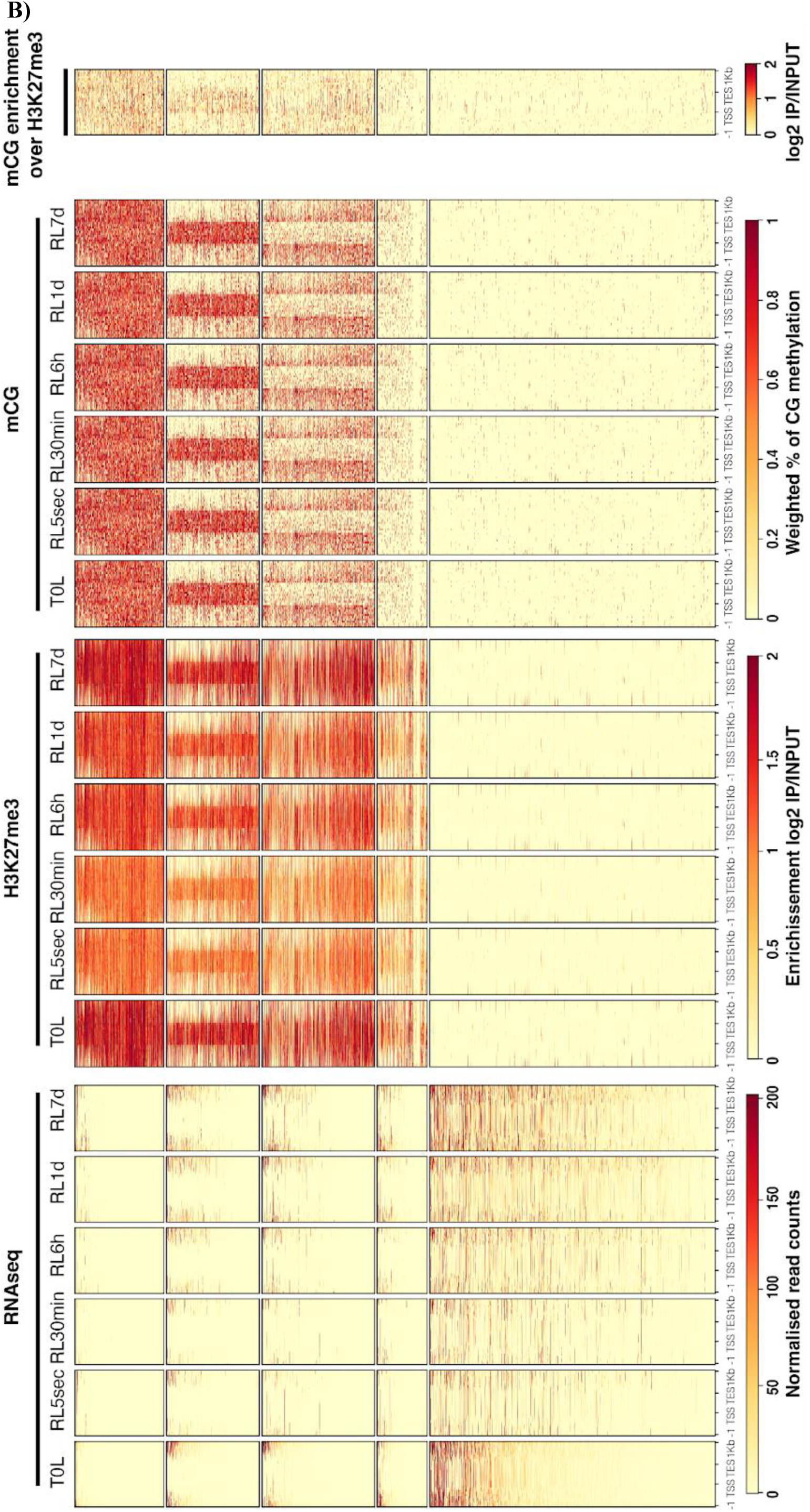
Co-occurrence of DNA methylation and H3K27me3 marks is linked to gene and TE silencing. **(A)** Heatmap displaying the distribution of RNA-seq reads, H3K27me3 (log₂ IP/INPUT), and CG methylation (% of mCG) over time at genes. (**B**) Same analysis for TEs larger than 100 bp. Cytosine methylation of H3K27me3-enriched chromatin fragments was confirmed experimentally through nanopore sequencing after H3K27me3 ChIP (see Methods).

Correlation of marking by DNA methylation and H3K27me3 in bulk analyses of cell populations does not demonstrate that both marks occur simultaneously on the same molecule/nucleosome, with distinct cells potentially bearing one or the other mark at a given locus. To test their potential co-occurrence *in situ*, we performed long-read ONT sequencing of DNA recovered after a H3K27me3 ChIP. This approach revealed a substantial increase in DNA methylation levels in the ChIP samples compared to input DNA (**Figure 5, Figure S10B**), thereby providing direct evidence that DNA fragments bound by H3K27me3-containing nucleosomes are enriched in DNA methylation. Although correlation marking between H3K27 and DNA methylation has emerged in groups like Stramenopiles, Ciliates, and Bryophytes (Veluchamy et al., 2015; Frapporti et al., 2019; Montgomery et al., 2020), this study provides the first demonstration of H3K27me3 and DNA methylation co-marking in diatoms.

## Discussion

By profiling the chromatin marks H2Bub, H3K27me3, and CG methylation, this study provides the first insights into chromatin dynamics in a polar microalga. During conditions mimicking extreme environmental fluctuations, the entry and exit of the polar night, we show that H2Bub levels change dynamically with gene expression in *F. cylindrus*. In contrast, H3K27me3 and DNA methylation, which repress mainly TEs and a few genes, remain globally stable. This stability underscores the necessity of maintaining chromatin organization and transcriptional control under prolonged darkness. Additionally, we show that H3K27me3 and DNA methylation frequently co-occur on the same genomic regions, presumably acting in concert to maintain silencing. While previous studies (Veluchamy et al., 2015; Frapporti et al., 2019; Montgomery et al., 2020) have hinted at this interplay, our findings provide the first evidence of H3K27me3 and DNA methylation co-marking in microalgae. This discovery opens new avenues for investigating the functional interactions between these chromatin modifications in the evolution of unicellular eukaryotes.

### H2Bub dynamics as a marker of transcriptional regulation in response to light cues in *F. cylindrus*

We observed that H2Bub enrichment at genes correlates with increased transcript levels, potentially modulating transcription during transitions between light and darkness as well as during light recovery. Notably, key survival genes remain regulated by H2Bub in the dark such as genes involved in BCAA, known to play a role in environmental adaptation in diatoms and could serve as an alternate energy source in darkness (Joli et al. 2024). In contrast, prior to darkness and upon re-exposure to light, H2Bub is enriched at genes associated with acclimation to continuous light conditions. This indicates that, beyond its role in transcriptional activation, H2Bub dynamically regulates condition-specific functions crucial for both dark survival and light adaptation, similar to its role in *A. thaliana* during the initial exposure to light post-germination (Bourbousse et al., 2012). The limited dynamics observed in *F. cylindrus* during darkness warrant further investigation. In *A. thaliana*, methods such as RNA spiking and ChIP-RNA Pol II have enabled the absolute quantification of transcript levels and the study of actively transcribed genes, refining our understanding of light-mediated gene regulation (Schivre et al., 2025). Applying these approaches to *F. cylindrus* could help link transcriptome dynamics with epigenomic data, shedding light on how cellular energy status and/or light signals drive regulatory changes and contribute to the previously suggested hypometabolic state (Joli et al., 2024). Our findings establish H2Bub as an evolutionarily conserved mark of transcriptional activation in Stramenopiles, and future functional studies, such as mutating homologous *Bre1* candidate genes, could offer further insights into its regulatory role.

### Open chromatin in prolonged darkness: a potential strategy for rapid acclimation to light re-exposure?

H3K27me3-marked genes and TEs remain repressed regardless of enrichment fluctuations nor light conditions. Among the few H3K27me3-marked genes that were expressed, no correlation was observed between transcript levels and H3K27me3 enrichment. This may reflect heterogeneity within the cell population, where a small subset of cells loses H3K27me3 rather than a uniform decrease across the culture. However, if this loss systematically triggered transcriptional activation, a stronger RNA-seq correlative signal would be expected, which was not the case. This suggests that occasional gene expression may result from stochastic H3K27me3 loss in a few cells rather than a regulated response. Single-cell approaches would be needed to distinguish between these possibilities.

Among the few expressed TEs marked by H3K27me3, we identified two that are located within gene introns. Notably, these TEs contain TF binding sites, a phenomenon also observed in H3K27me3-marked TEs in bryophytes and *Arabidopsis*. It has been proposed that TEs are repurposed as cis-elements, binding specific TF to regulate gene expression (Hisanaga et al., 2023). These findings highlight the need for further investigation into the regulatory roles of TEs in transcriptional networks. Many TEs that are not marked by H3K27me3 appear to be expressed in our experiment. Upon visual inspection of some candidates we found that RNA-seq reads are not confined to the body of the TE but extend upstream and downstream, with unannotated genes or genes located nearby. A more thorough manual curation of TE annotations would be necessary to confirm the final list of TEs in *F. cylindrus*.

In this study, we did not detect an expansion of H3K27me3 to new loci during prolonged darkness. This suggests that facultative heterochromatin is not the primary mechanism behind the extensive gene downregulation associated with hypometabolism in the dark. While PRC2 is involved in key regulatory processes such as cell differentiation in *P. tricornutum* (Zhao et al., 2021; Zarif et al., 2024) and thermo-morphogenesis in *A. thaliana* (Kim et al., 2023), its stability in *F. cylindrus* may reflect an adaptive strategy. We hypothesize that the light-dark transitions in our experimental setup likely do not affect PRC2 or do not constitute a significant environmental stress for this polar diatom, which is naturally acclimated to extreme light fluctuations. Although previous studies indicate an increase in the heterochromatin-to-euchromatin ratio during prolonged darkness in this species (Joli et al., 2024), our findings suggest that an open chromatin state may be maintained at many genes under prolonged darkness, possibly facilitating rapid cellular adaptations to the return of stressful light exposure.

### Dual-Locking Mechanism: Co-occurrence of H3K27me3 and DNA methylation

The mirroring profiles of DNA methylation and H3K27me3 across genes and TEs suggest a coordinated role in chromatin regulation in *F. cylindrus*. Studies in Stramenopiles, Ciliates, and Bryophytes indicate that H3K27me3 often serves as the dominant silencing mark on TEs, frequently co-occurring with H3K9me3 or DNA methylation (Veluchamy et al., 2015; Frapporti et al., 2019; Montgomery et al., 2020).

Using nanopore sequencing of H3K27me3-immunoprecipitated DNA, we confirmed that both marks are present on the same DNA fragments. This finding suggests a potential synergy between these two silencing systems, forming a dual-locking mechanism that ensures stable repression. This supports the hypothesis that one of the ancestral functions of the PRC2 was to defend against intragenomic parasites, such as TEs, before being co-opted for developmental roles in multicellular eukaryotes (Deleris et al. 2021). Further exploration, especially of H3K9me3 profiles, requires a dedicated analysis by ChIP-seq However, the absence of sufficiently specific H3K9me3 antibodies for *F. cylindrus* limited this approach.

The slight dynamics of H3K27me3, compared to the stability of DNA methylation, raise intriguing questions about their respective roles. Beyond potential heterogeneity within the cell population, these two marks may serve distinct functions: H3K27me3 could act as a dynamic, adaptive signal, while DNA methylation might function as a more stable epigenetic memory, facilitating the restoration of H3K27me3 upon light re-exposure. This raises an important question: what is the remaining level of H3K27me3 after a longer period of darkness (polar night lasting 6 months at the pole)? Would the proposed dual-locking mechanism become necessary to prevent the activation of TEs under even more drastic conditions? Additionally, among the genes marked by H3K27me3, we identified a potential homolog of HP1, a known reader of H3K9me, as well as genes involved in chromatin organization and regulation. In contrast, H2Bub enrichment was predominantly observed at genes involved in cellular metabolism, highlighting the distinct roles of these marks. A complete loss of H3K27me3 at TEs and these genes could disrupt chromatin architecture and genome stability during the polar night, and the stability of DNA methylation suggests that maintaining repression of TEs is a crucial adaptive strategy. Thus, the dual-locking mechanism may provide an evolutionary advantage by ensuring the stable repression of these elements, thereby preserving genome integrity despite fluctuating environmental conditions.

DNA methylation has been associated with gene and TEs repression in various organisms, including plants, fungi, and vertebrates (Law and Jacobsen, 2010; Zemach and Zilberman, 2010). In *P. tricornutum*, for example, extensive methylation regulates nitrate-responsive genes (Veluchamy et al., 2013). However, exceptions to this trend, where DNA methylation does not correlate with gene expression, have also been observed (Huff and Zilberman, 2014). These inconsistencies highlight the complexity of DNA methylation’s roles in regulating gene expression. In our experiment, the lack of significant changes in DNA methylation and H3K27me3 during darkness or light recovery suggests these marks are not key to transcriptional changes in *F. cylindrus*.

### Toward a deeper deciphering of the epigenomic code in *F. cylindrus*?

The analysis of H3K27me3 and H2Bub landscapes revealed a functional dichotomy between these two chromatin marks. H3K27me3 is largely absent from genes enriched in H2Bub, suggesting a spatial segregation of activation (H2Bub) and repression (H3K27me3) marks. This opposing distribution points to a layered regulatory mechanism where these marks may act antagonistically to fine-tune transcriptional output. Notably, genes marked by H3K27me3 at upstream regions exhibited a higher frequency of expression compared to those marked within their body but a lower transcript level than those unmarked, illustrating the complexity of chromatin-based regulation in *F. cylindrus*. A similar observation was made in *P. tricornutum*, where it was proposed that these upstream or downstream regions might act as bivalent domains, harboring additional marks that could modulate gene expression in a context-dependent manner (Veluchamy et al., 2015).

The global transcriptional repression observed in darkness does not appear to be directly linked to specific H3K27me3 or DNA methylation patterns. Likewise, some genes and TEs remain permanently silent despite lacking detectable levels of these marks. This suggests the involvement of alternative silencing mechanisms, an intrinsic ability of certain genes to remain inactive without active repression, or the targeted removal of transcription-associated chromatin marks, such as histone deacetylation. The role of histone acetylation and other marks associated with transcriptional activation or constitutive heterochromatin remains largely unexplored in diatoms. A key limitation of our study is that population-level analyses may obscure heterogeneity in chromatin states across individual cells. While we expect a degree of synchronization among cells exposed to darkness, future single-cell approaches could offer a more precise understanding of chromatin regulation and transcriptional responses in this polar diatom.

## References

Akalin, A., Kormaksson, M., Li, S., Garrett-Bakelman, F.E., Figueroa, M.E., Melnick, A. and Mason, C.E., 2012. methylKit: a comprehensive R package for the analysis of genome-wide DNA methylation profiles. Genome biology, 13, pp.1–9.

Bolger, A.M., Lohse, M. and Usadel, B., 2014. Trimmomatic: a flexible trimmer for Illumina sequence data. Bioinformatics, 30(15), pp.2114–2120.

Bourbousse, C., Ahmed, I., Roudier, F., Zabulon, G., Blondet, E., Balzergue, S., Colot, V., Bowler, C. and Barneche, F., 2012. Histone H2B monoubiquitination facilitates the rapid modulation of gene expression during Arabidopsis photomorphogenesis. PLoS genetics, 8(7), p.e1002825.

Bourbousse, C., Mestiri, I., Zabulon, G., Bourge, M., Formiggini, F., Koini, M.A., Brown, S.C., Fransz, P., Bowler, C. and Barneche, F., 2015. Light signaling controls nuclear architecture reorganization during seedling establishment. Proceedings of the National Academy of Sciences, 112(21), pp.E2836–E2844.

Boyer, L.A., Plath, K., Zeitlinger, J., Brambrink, T., Medeiros, L.A., Lee, T.I., Levine, S.S., Wernig, M., Tajonar, A., Ray, M.K. and Bell, G.W., 2006. Polycomb complexes repress developmental regulators in murine embryonic stem cells. nature, 441(7091), pp.349–353.

Chittock, E.C., Latwiel, S., Miller, T.C. and Müller, C.W., 2017. Molecular architecture of polycomb repressive complexes. Biochemical Society Transactions, 45(1), pp.193–205.

Deleris, A., Berger, F. and Duharcourt, S., 2021. Role of Polycomb in the control of transposable elements. Trends in Genetics, 37(10), pp.882–889.

Dumesic, P.A., Homer, C.M., Moresco, J.J., Pack, L.R., Shanle, E.K., Coyle, S.M., Strahl, B.D., Fujimori, D.G., Yates, J.R. and Madhani, H.D., 2015. Product binding enforces the genomic specificity of a yeast polycomb repressive complex. Cell, 160(1), pp.204–218.

Falciatore, A., Jaubert, M., Bouly, J.P., Bailleul, B. and Mock, T., 2020. Diatom molecular research comes of age: model species for studying phytoplankton biology and diversity. The Plant Cell, 32(3), pp.547–572.

Field, C.B., Behrenfeld, M.J., Randerson, J.T. and Falkowski, P., 1998. Primary production of the biosphere: integrating terrestrial and oceanic components. science, 281(5374), pp.237–240.

Filion, G.J., van Bemmel, J.G., Braunschweig, U., Talhout, W., Kind, J., Ward, L.D., Brugman, W., de Castro, I.J., Kerkhoven, R.M., Bussemaker, H.J. and van Steensel, B., 2010. Systematic protein location mapping reveals five principal chromatin types in Drosophila cells. Cell, 143(2), pp.212–224.

Flutre, T., Duprat, E., Feuillet, C. and Quesneville, H., 2011. Considering transposable element diversification in de novo annotation approaches. PloS one, 6(1), p.e16526.

Frapporti, A., Miró Pina, C., Arnaiz, O., Holoch, D., Kawaguchi, T., Humbert, A., Eleftheriou, E., Lombard, B., Loew, D., Sperling, L. and Guitot, K., 2019. The Polycomb protein Ezl1 mediates H3K9 and H3K27 methylation to repress transposable elements in Paramecium. Nature communications, 10(1), p.2710.

Fuchs, G., Hollander, D., Voichek, Y., Ast, G. and Oren, M., 2014. Cotranscriptional histone H2B monoubiquitylation is tightly coupled with RNA polymerase II elongation rate. Genome research, 24(10), pp.1572–1583.

Grasser, K.D., Rubio, V. and Barneche, F., 2021. Multifaceted activities of the plant SAGA complex. Biochimica et Biophysica Acta (BBA)-Gene Regulatory Mechanisms, 1864(2), p.194613.

Henry, K.W., Wyce, A., Lo, W.S., Duggan, L.J., Emre, N.T., Kao, C.F., Pillus, L., Shilatifard, A., Osley, M.A. and Berger, S.L., 2003. Transcriptional activation via sequential histone H2B ubiquitylation and deubiquitylation, mediated by SAGA-associated Ubp8. Genes & development, 17(21), pp.2648–2663.

Hisanaga, T., Romani, F., Wu, S., Kowar, T., Wu, Y., Lintermann, R., Fridrich, A., Cho, C.H., Chaumier, T., Jamge, B. and Montgomery, S.A., 2023. The Polycomb repressive complex 2 deposits H3K27me3 and represses transposable elements in a broad range of eukaryotes. Current Biology, 33(20), pp.4367–4380

Hoguin, A., Yang, F., Groisillier, A., Bowler, C., Genovesio, A., Ait-Mohamed, O., Vieira, F.R.J. and Tirichine, L., 2023. The model diatom Phaeodactylum tricornutum provides insights into the diversity and function of microeukaryotic DNA methyltransferases. Communications Biology, 6(1), p.253.

Huff, J.T. and Zilberman, D., 2014. Dnmt1-independent CG methylation contributes to nucleosome positioning in diverse eukaryotes. Cell, 156(6), pp.1286–1297.

Ibarbalz, F.M., Henry, N., Brandão, M.C., Martini, S., Busseni, G., Byrne, H., Coelho, L.P., Endo, H., Gasol, J.M., Gregory, A.C. and Mahé, F., 2019. Global trends in marine plankton diversity across kingdoms of life. Cell, 179(5), pp.1084–1097.

Jamieson, K., Rountree, M.R., Lewis, Z.A., Stajich, J.E. and Selker, E.U., 2013. Regional control of histone H3 lysine 27 methylation in Neurospora. Proceedings of the National Academy of Sciences, 110(15), pp.6027–6032.

Jin, Y., Tam, O.H., Paniagua, E. and Hammell, M., 2015. TEtranscripts: a package for including transposable elements in differential expression analysis of RNA-seq datasets. Bioinformatics, 31(22), pp.3593–3599.

Joli, N., Concia, L., Mocaer, K., Guterman, J., Laude, J., Guerin, S., Sciandra, T., Bruyant, F., Ait-Mohamed, O., Beguin, M. and Forget, M.H., 2024. Hypometabolism to survive the long polar night and subsequent successful return to light in the diatom Fragilariopsis cylindrus. New Phytologist, 241(5), pp.2193–2208.

Kato, M., Miura, A., Bender, J., Jacobsen, S.E. and Kakutani, T., 2003. Role of CG and non-CG methylation in immobilization of transposons in Arabidopsis. Current Biology, 13(5), pp.421–426.

Kharchenko, P.V., Alekseyenko, A.A., Schwartz, Y.B., Minoda, A., Riddle, N.C., Ernst, J., Sabo, P.J., Larschan, E., Gorchakov, A.A., Gu, T. and Linder-Basso, D., 2011. Comprehensive analysis of the chromatin landscape in Drosophila melanogaster. Nature, 471(7339), pp.480–485.

Kim, J., Bordiya, Y., Xi, Y., Zhao, B., Kim, D.H., Pyo, Y., Zong, W., Ricci, W.A. and Sung, S., 2023. Warm temperature-triggered developmental reprogramming requires VIL1-mediated, genome-wide H3K27me3 accumulation in Arabidopsis. Development, 150(5), p.dev201343.

Krueger, F. and Andrews, S.R., 2011. Bismark: a flexible aligner and methylation caller for Bisulfite-Seq applications. bioinformatics, 27(11), pp.1571–1572.

Law, J.A. and Jacobsen, S.E., 2010. Establishing, maintaining and modifying DNA methylation patterns in plants and animals. Nature Reviews Genetics, 11(3), pp.204–220.

Lin, X., Tirichine, L. and Bowler, C., 2012. Protocol: Chromatin immunoprecipitation (ChIP) methodology to investigate histone modifications in two model diatom species. Plant methods, 8, pp.1–9.

Margueron, R. and Reinberg, D., 2011. The Polycomb complex PRC2 and its mark in life. Nature, 469(7330), pp.343–349.

Marsano, R.M., Giordano, E., Messina, G. and Dimitri, P., 2019. A new portrait of constitutive heterochromatin: lessons from Drosophila melanogaster. Trends in Genetics, 35(9), pp.615–631.

Maumus, F., Allen, A.E., Mhiri, C., Hu, H., Jabbari, K., Vardi, A., Grandbastien, M.A. and Bowler, C., 2009. Potential impact of stress activated retrotransposons on genome evolution in a marine diatom. BMC genomics, 10, pp.1–19.

Minsky, N., Shema, E., Field, Y., Schuster, M., Segal, E. and Oren, M., 2008. Monoubiquitinated H2B is associated with the transcribed region of highly expressed genes in human cells. Nature cell biology, 10(4), pp.483–488.

Mock, T., Otillar, R.P., Strauss, J., McMullan, M., Paajanen, P., Schmutz, J., Salamov, A., Sanges, R., Toseland, A., Ward, B.J. and Allen, A.E., 2017. Evolutionary genomics of the cold-adapted diatom Fragilariopsis cylindrus. Nature, 541(7638), pp.536–540.

Montgomery, S.A., Tanizawa, Y., Galik, B., Wang, N., Ito, T., Mochizuki, T., Akimcheva, S., Bowman, J.L., Cognat, V., Maréchal-Drouard, L. and Ekker, H., 2020. Chromatin organization in early land plants reveals an ancestral association between H3K27me3, transposons, and constitutive heterochromatin. Current Biology, 30(4), pp.573–588.

Nassrallah, A., Rougée, M., Bourbousse, C., Drevensek, S., Fonseca, S., Iniesto, E., Ait-Mohamed, O., Deton-Cabanillas, A.F., Zabulon, G., Ahmed, I. and Stroebel, D., 2018. DET1-mediated degradation of a SAGA-like deubiquitination module controls H2Bub homeostasis. Elife, 7, p.e37892.

Ni, P., Huang, N., Nie, F., Zhang, J., Zhang, Z., Wu, B., Bai, L., Liu, W., Xiao, C.L., Luo, F. and Wang, J., 2021. Genome-wide detection of cytosine methylations in plant from Nanopore data using deep learning. Nature communications, 12(1), p.5976.

Oss-Ronen, L., Sarusi, T. and Cohen, I., 2022. Histone mono-ubiquitination in transcriptional regulation and its mark on life: emerging roles in tissue development and disease. Cells, 11(15), p.2404.

Pauler, F.M., Sloane, M.A., Huang, R., Regha, K., Koerner, M.V., Tamir, I., Sommer, A., Aszodi, A., Jenuwein, T. and Barlow, D.P., 2009. H3K27me3 forms BLOCs over silent genes and intergenic regions and specifies a histone banding pattern on a mouse autosomal chromosome. Genome research, 19(2), pp.221–233.

Pierella Karlusich, J.J., Ibarbalz, F.M. and Bowler, C., 2020. Phytoplankton in the Tara ocean. Annual Review of Marine Science, 12(1), pp.233–265.

Quesneville, H., Bergman, C.M., Andrieu, O., Autard, D., Nouaud, D., Ashburner, M. and Anxolabehere, D., 2005. Combined evidence annotation of transposable elements in genome sequences. PLoS computational biology, 1(2), p.e22.

Rice, P., Longden, I. and Bleasby, A., 2000. EMBOSS: the European molecular biology open software suite. Trends in genetics, 16(6), pp.276–277.

Roudier, F., Ahmed, I., Bérard, C., Sarazin, A., Mary-Huard, T., Cortijo, S., Bouyer, D., Caillieux, E., Duvernois-Berthet, E., Al-Shikhley, L. and Giraut, L., …Colot, V. 2011. Integrative epigenomic mapping defines four main chromatin states in Arabidopsis. The EMBO journal, 30(10), pp.1928–1938.

Schivre, G., Wolff, L., Mirasole, F.M., Armanet, E., Davidson, M.L., Vidal, A., Cuménal, D., Dumont, M., Bourge, M., Baroux, C. and Bourbousse, C., 2025. Genome-scale transcriptome augmentation during Arabidopsis thaliana photomorphogenesis. bioRxiv, pp.2025–01

Schubert, D., Clarenz, O. and Goodrich, J., 2005. Epigenetic control of plant development by Polycomb-group proteins. Current opinion in plant biology, 8(5), pp.553–561.

Schuettengruber, B., Bourbon, H.M., Di Croce, L. and Cavalli, G., 2017. Genome regulation by polycomb and trithorax: 70 years and counting. Cell, 171(1), pp.34–57.

Sequeira-Mendes, J., Aragüez, I., Peiró, R., Mendez-Giraldez, R., Zhang, X., Jacobsen, S.E., Bastolla, U. and Gutierrez, C., 2014. The functional topography of the Arabidopsis genome is organized in a reduced number of linear motifs of chromatin states. The Plant Cell, 26(6), pp.2351–2366.

Sharaf, A., Vijayanathan, M., Oborník, M. and Mozgová, I., 2022. Phylogenetic profiling resolves early emergence of PRC2 and illuminates its functional core. Life Science Alliance, 5(7).

Shaver, S., Casas-Mollano, J.A., Cerny, R.L. and Cerutti, H., 2010. Origin of the polycomb repressive complex 2 and gene silencing by an E (z) homolog in the unicellular alga Chlamydomonas. Epigenetics, 5(4), pp.301–312.

Supek, F., Bošnjak, M., Škunca, N. and Šmuc, T., 2011. REVIGO summarizes and visualizes long lists of gene ontology terms. PloS one, 6(7), p.e21800.

Tanny, J.C., Erdjument-Bromage, H., Tempst, P. and Allis, C.D., 2007. Ubiquitylation of histone H2B controls RNA polymerase II transcription elongation independently of histone H3 methylation. Genes & development, 21(7), pp.835–847.

Veluchamy, A., Lin, X., Maumus, F., Rivarola, M., Bhavsar, J., Creasy, T., O’Brien, K., Sengamalay, N.A., Tallon, L.J., Smith, A.D. and Rayko, E., 2013. Insights into the role of DNA methylation in diatoms by genome-wide profiling in Phaeodactylum tricornutum. Nature communications, 4(1), p.2091.

Veluchamy, A., Rastogi, A., Lin, X., Lombard, B., Murik, O., Thomas, Y., Dingli, F., Rivarola, M., Ott, S., Liu, X. and Sun, Y., 2015. An integrative analysis of post-translational histone modifications in the marine diatom Phaeodactylum tricornutum. Genome biology, 16, pp.1–18.

Vijayanathan, M., Trejo-Arellano, M.G. and Mozgová, I., 2022. Polycomb repressive complex 2 in eukaryotes—an evolutionary perspective. Epigenomes, 6(1), p.3.

Walworth, N.G., Lee, M.D., Dolzhenko, E., Fu, F.X., Smith, A.D., Webb, E.A. and Hutchins, D.A., 2021. Long-term m5C methylome dynamics parallel phenotypic adaptation in the cyanobacterium Trichodesmium. Molecular Biology and Evolution, 38(3), pp.927–939

Zarif, M., Rousselot, E., Jesus, B., Tirichine, L. and Duc, C., 2024. H3K27me3 and EZH Are Involved in the Control of the Heat-Stress-Elicited Morphological Changes in Diatoms. International Journal of Molecular Sciences, 25(15), p.8373

Zemach, A. and Zilberman, D., 2010. Evolution of eukaryotic DNA methylation and the pursuit of safer sex. Current Biology, 20(17), pp.R780–R785.

Zentner, G.E. and Henikoff, S., 2013. Regulation of nucleosome dynamics by histone modifications. Nature structural & molecular biology, 20(3), pp.259–266.

Zhang, X., Clarenz, O., Cokus, S., Bernatavichute, Y.V., Pellegrini, M., Goodrich, J. and Jacobsen, S.E., 2007. Whole-genome analysis of histone H3 lysine 27 trimethylation in Arabidopsis. PLoS biology, 5(5), p.e129.

Zhao, X., Rastogi, A., Deton Cabanillas, A.F., Ait Mohamed, O., Cantrel, C., Lombard, B., Murik, O., Genovesio, A., Bowler, C., Bouyer, D. and Loew, D., 2021. Genome wide natural variation of H3K27me3 selectively marks genes predicted to be important for cell differentiation in Phaeodactylum tricornutum. New Phytologist, 229(6), pp.3208–3220.

Zhao, X., Xiong, J., Mao, F., Sheng, Y., Chen, X., Feng, L., Dui, W., Yang, W., Kapusta, A., Feschotte, C. and Coyne, R.S., 2019. RNAi-dependent Polycomb repression controls transposable elements in Tetrahymena. Genes & Development, 33(5-6), pp.348–364.

