## Supplementary Figures for "Entry and exit of *Fragilariopsis cylindrus* diatoms from the polar night: evidence for chromatin-mediated genome control"

**from**

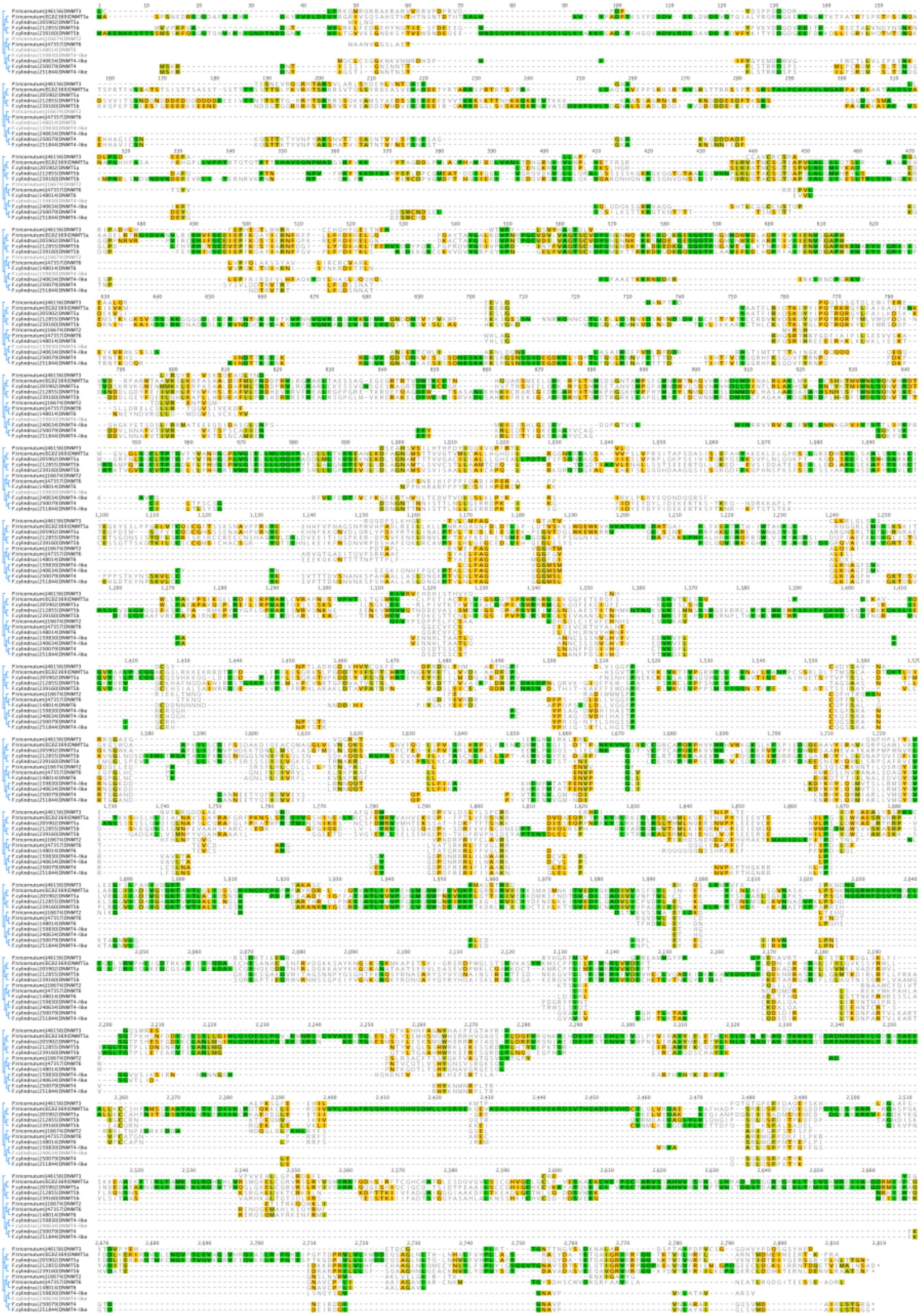

**Figure S1:** Alignment of DNMT candidates obtained from Hoguin et al. (2023) and KEGG Orthology. The left tree was built using the neighbor-joining method in Geneious. Colors indicate sequence identity: green (100%), brown (80–100%), yellow (60–80%), and gray (<60%).

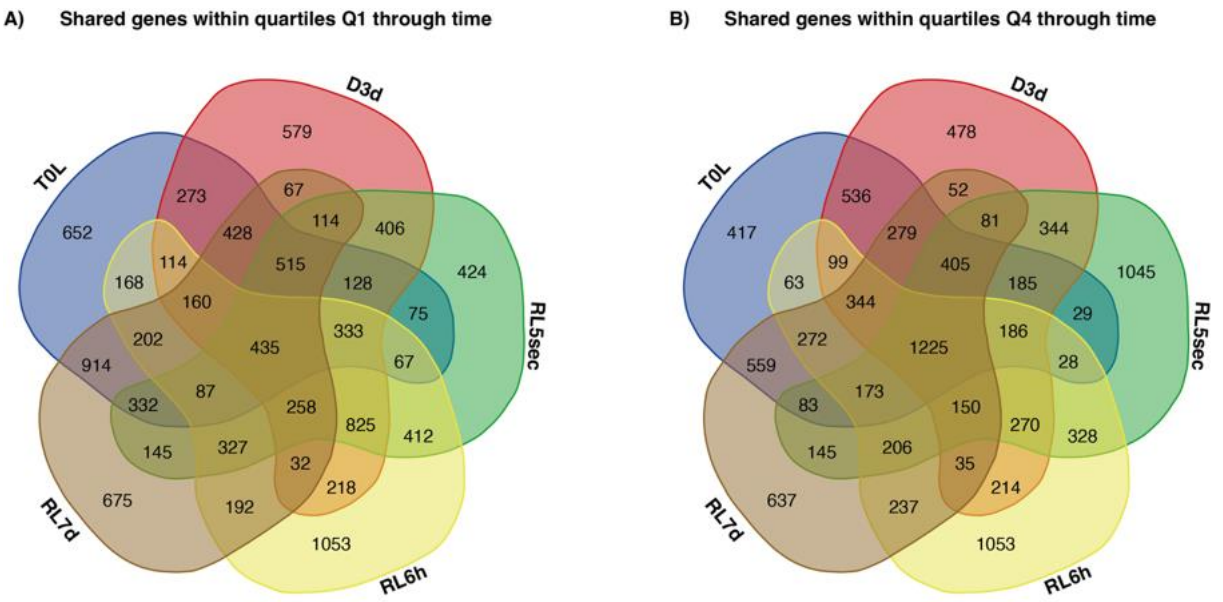


**Figure S2:** Venn diagram showing the intersection of genes across time points within expression quartiles from Figure 2B: **(A)** Q1 – top 25% least expressed genes and **(B)** Q4 – top 25% most highly expressed genes. Note that the sample labeled RL5sec refers to samples that were exposed to three months of darkness, followed by a brief 5-second flash of light.

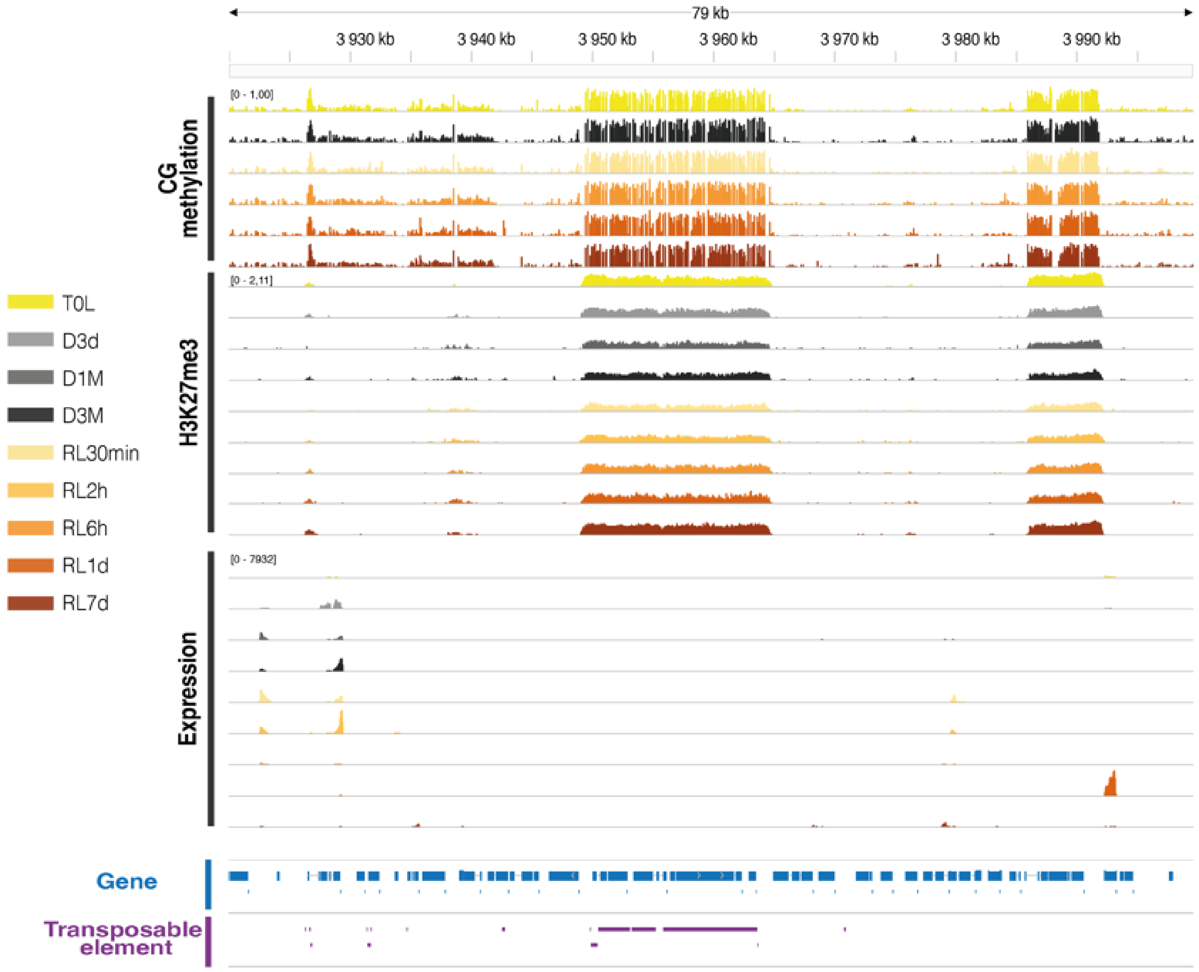


**Figure S3 :** Genome browser vizualisation of CG methylation, H3K27me3, and gene expression in relation to gene and TE annotations, highlighting the extensive H3K27me3 domains spanning several kilobases.

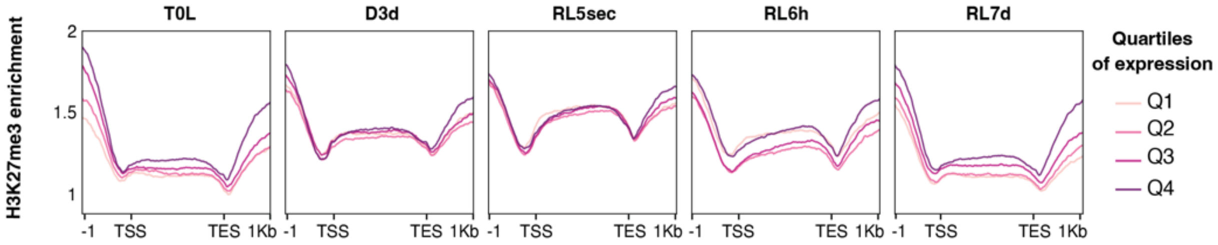


**Figure S4:** Metaplots of H3K27me3 enrichment (IP) across the same quartiles of expression illustrated in Figure 2.

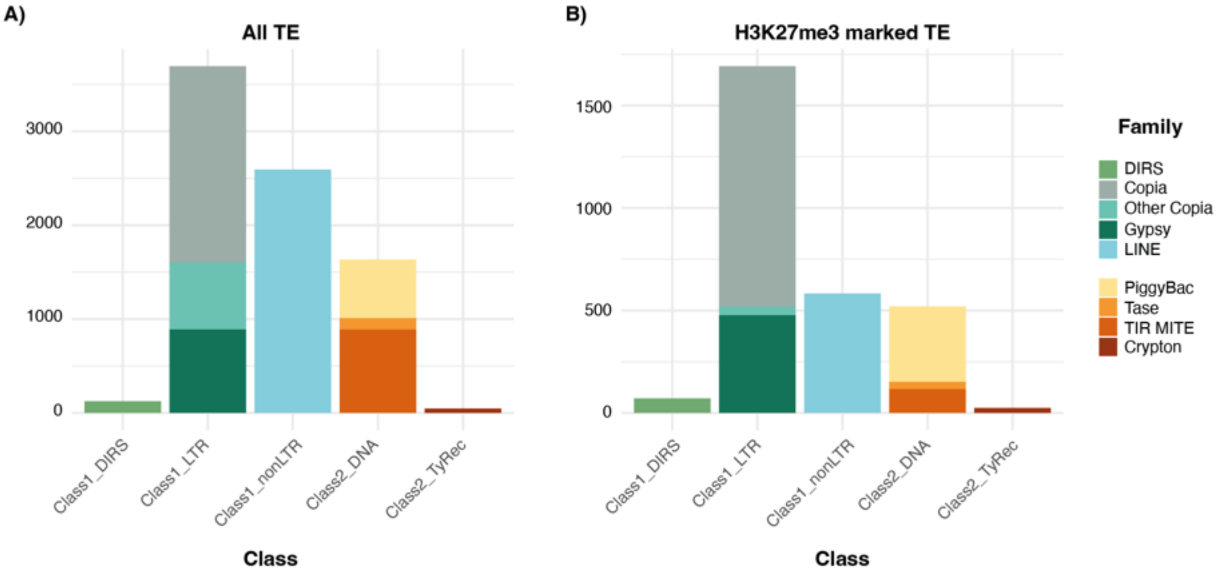


**Figure S5** : Distribution of functionally annotated TEs in *F. cylindrus* across different TE classes and families: (**A**) all TEs; (**B**) TEs marked by H3K27me3 at least at one time point.

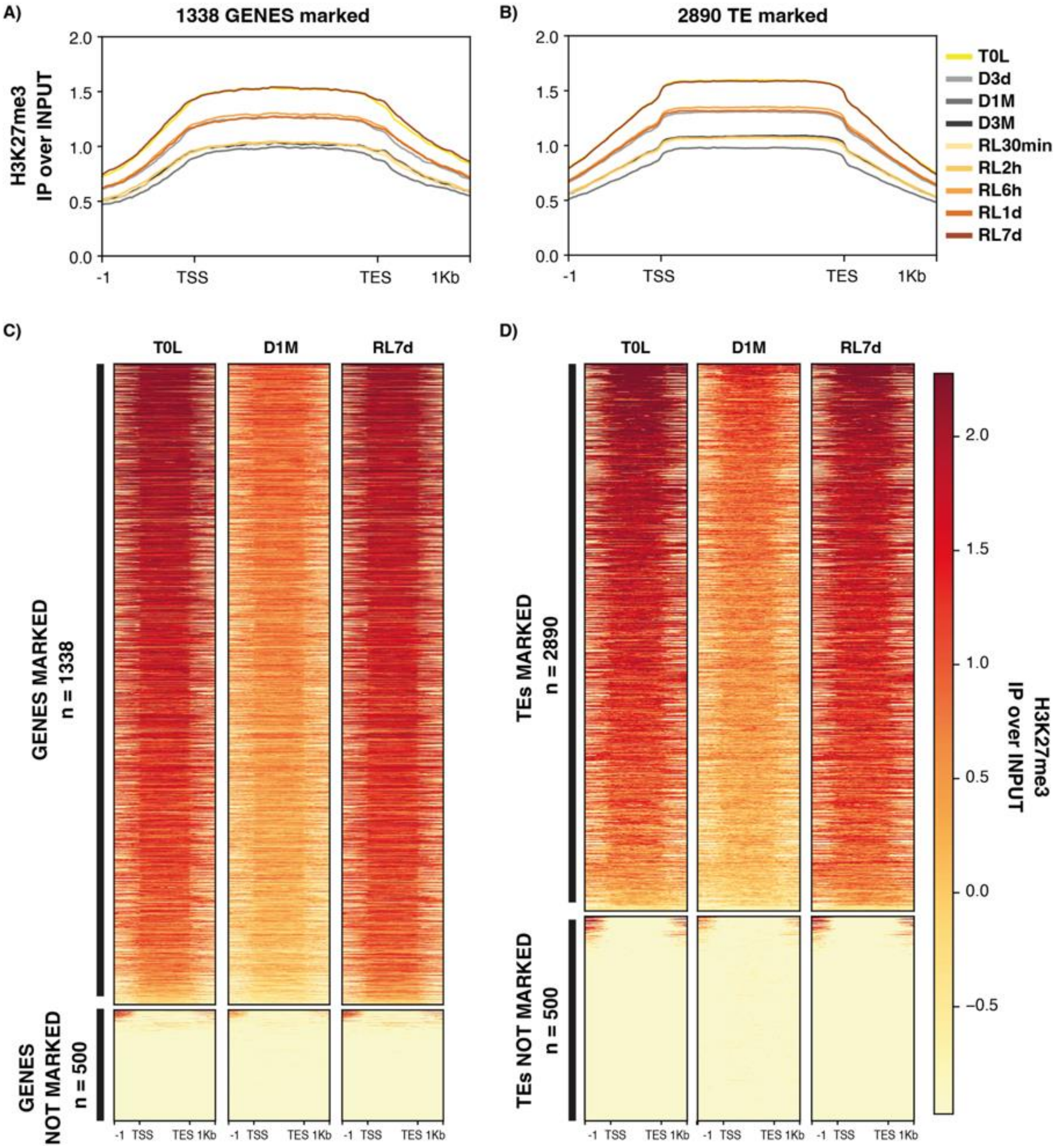


**Figure S6 : H3K27me3 is maintained over TEs and genes under prolonged darkness and subsequent light return.** (**A**) Metaplot of $\log_2$(IP/Input) for H3K27me3 at genes across time points. (**B**) Metaplot of $\log_2$(IP/Input) for H3K27me3 at transposable elements (TEs) across time points. (**C**) Heatmap of $\log_2$(IP/Input) for H3K27me3, showing consistently marked and unmarked genes. (**D**) Heatmap of $\log_2$(IP/Input) for H3K27me3, showing consistently marked and unmarked TEs. For clarity, only T0L (initial acclimation to full light), D1M (1 month of darkness), and RL7d (7 days after light re-exposure) are shown in the heatmaps. A random subset of 500 unmarked genes and TEs is also included.

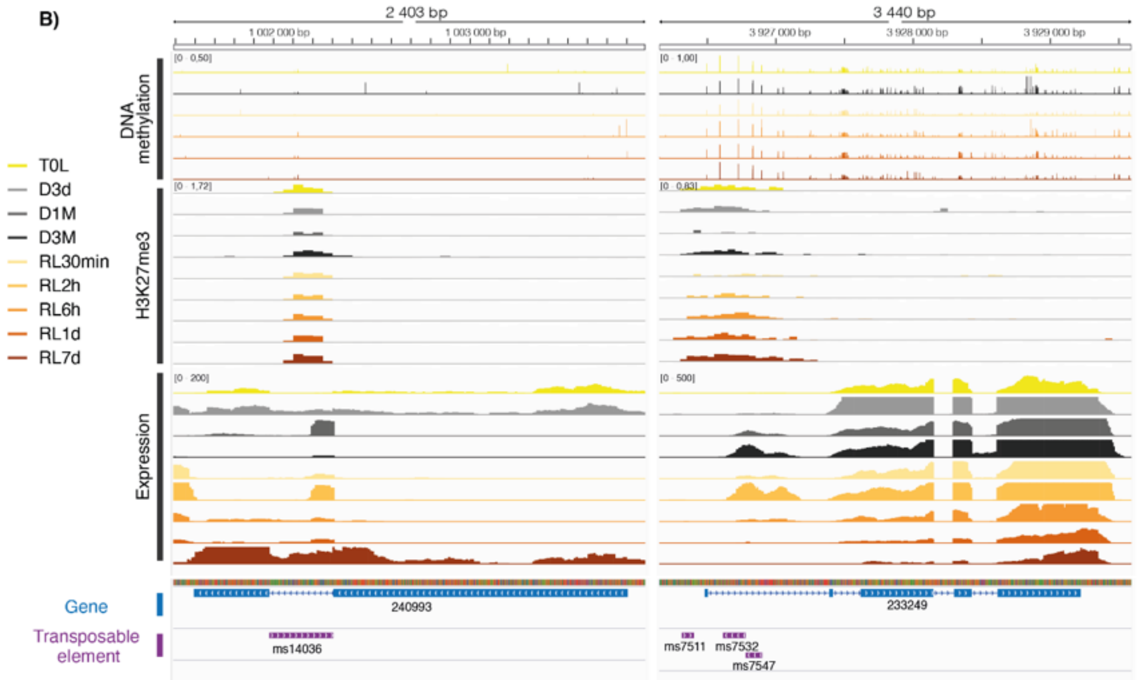


**Figure S7 :**: IGV screenshot showing the genomic context of the two TEs marked by H3K27me3 and having detectable read counts overlap with gene annotations, within intronic regions.

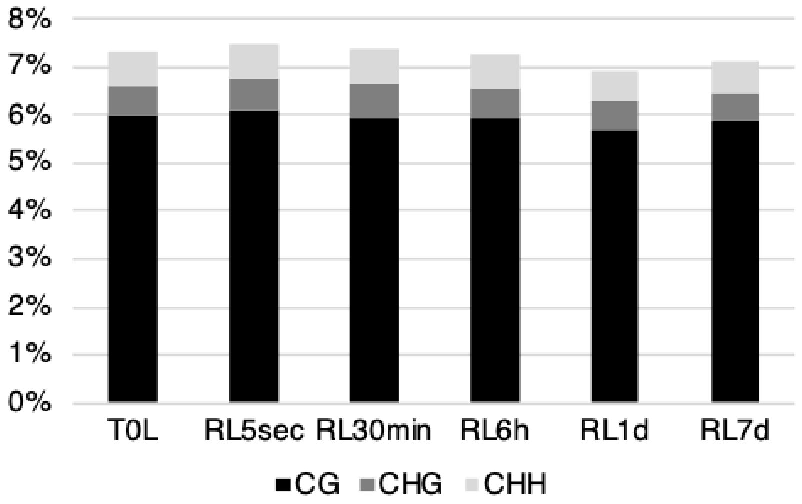


**Figure S8** : Percentage of genome coverage by DNA methylation in CG, CHG, and CHH contexts. The majority of DNA methylation occurs in the CG context (>80%), while CHG and CHH contexts show lower coverage.

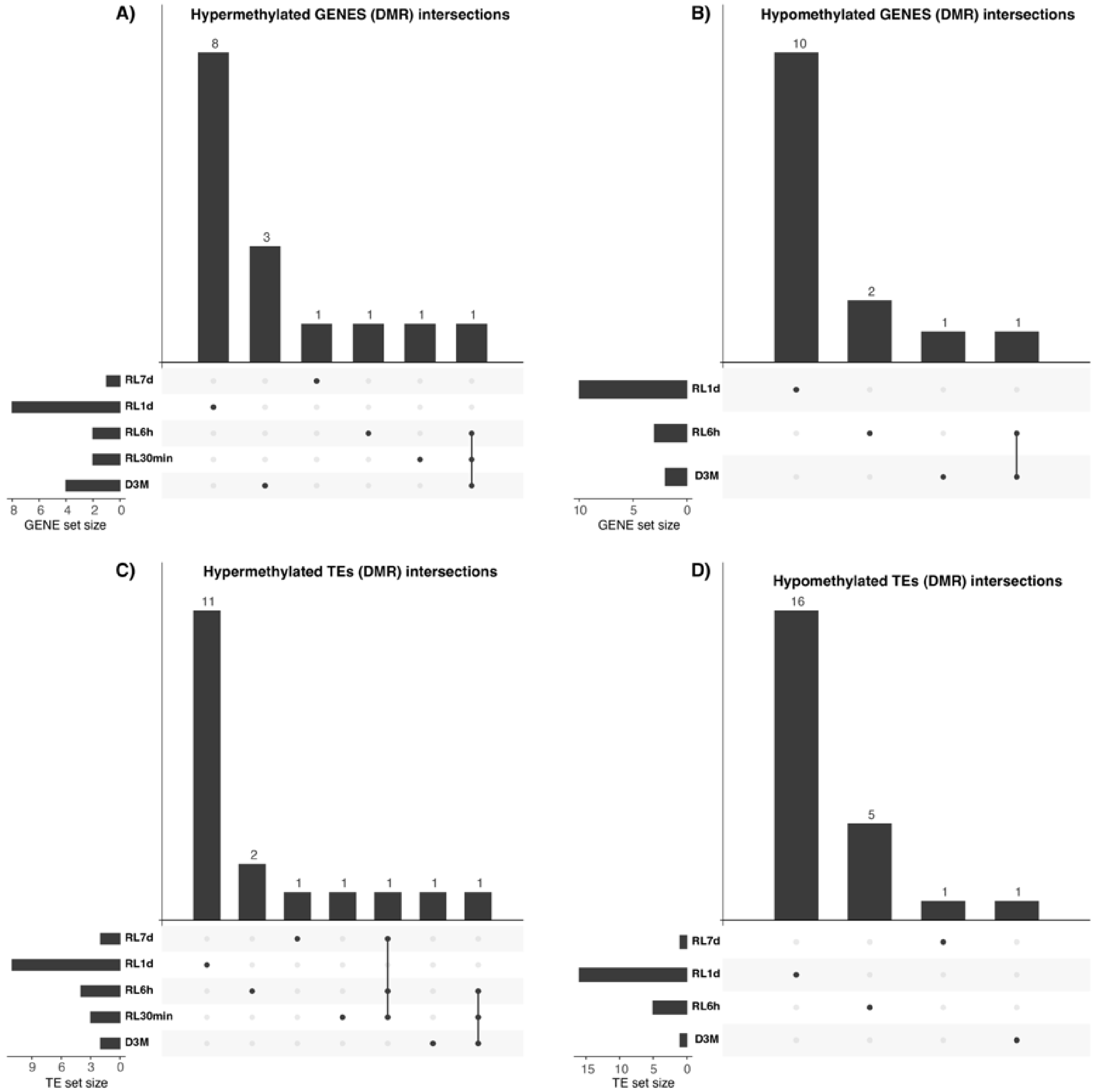


**Figure S9** : UpSet plots of shared differentially (**A**) hypermethylated genes (**B**) hypomethylated genes (**C**) hypermethylated TEs (**D**) hypomethylated TEs across time points.

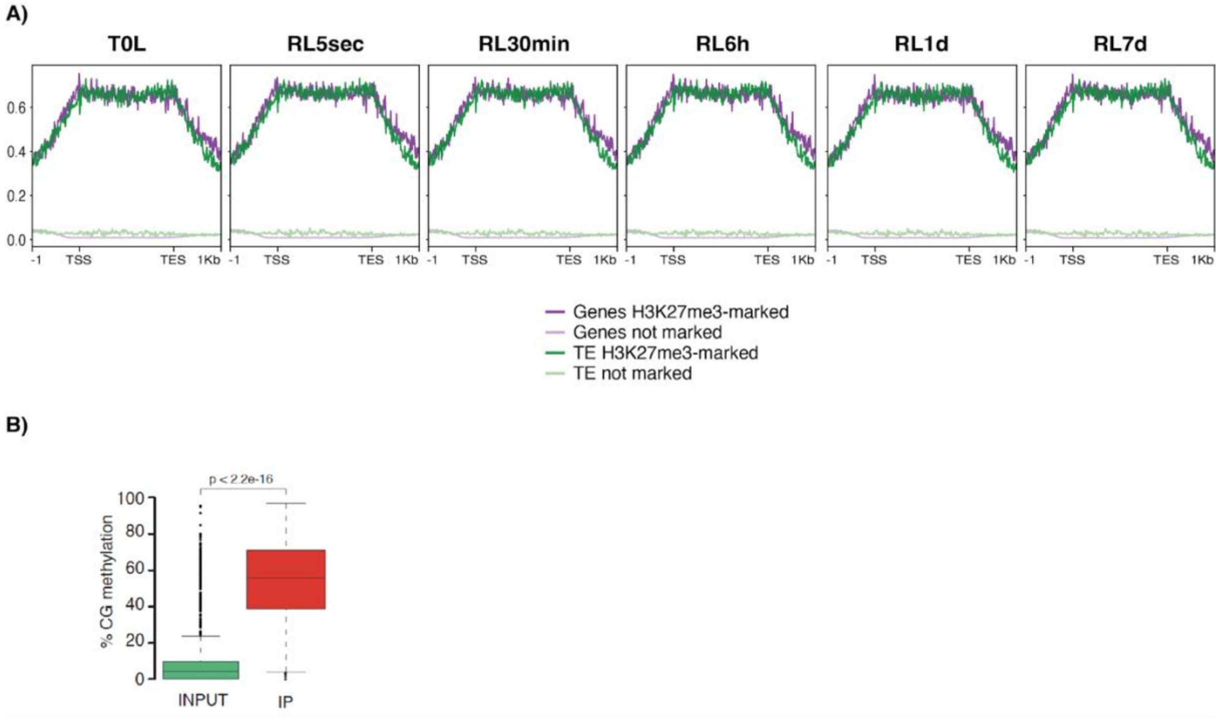


**Figure S10:** (**A**) Metaplot of DNA methylation from bisulfite sequencing over H3K27me3 marked and not marked genes and TEs over time. (**B**) Boxplot of the percentage of CG methylation in H3K27me3-immunoprecipitated DNA fragments compared to input DNA.
